# Redox-dependent synaptic clustering of gephyrin

**DOI:** 10.1101/2024.03.15.585209

**Authors:** Maria-Theresa Gehling, Filip Liebsch, Lianne Jacobs, Jan Riemer, Günter Schwarz

**Affiliations:** Institute of Biochemistry, Department of Chemistry, University of Cologne, 50674 Cologne, Germany; Cologne Excellence Cluster on Cellular Stress Responses in Aging-Associated Diseases (CECAD), University of Cologne, Cologne, Germany; Center for Molecular Medicine Cologne (CMMC), University of Cologne, Cologne, Germany

**Author notes:** Correspondence: Guenter Schwarz.

## Abstract

Reactive oxygen species (ROS) play a central role in enhancing inhibitory signal transmission, thus extending their role beyond oxidative stress in disease and aging. However, the underlying molecular mechanisms mediating these functions have remained elusive. At inhibitory synapses, the scaffolding protein gephyrin clusters glycine and GABA type A receptors. Since gephyrin harbors multiple surface-exposed cysteines, we investigated the regulatory influence of ROS on gephyrin. We show that H_2_O_2_-induced oxidation of gephyrin cysteines triggered reversible, synaptic multimerization through disulfide bridge formation, which provided more receptor binding sites, lead to proteolytic protection and enhanced liquid-liquid phase separation. We identified mitochondria-derived ROS as a physiological source and observed oxidized gephyrin multimers *in vivo,* indicating that gephyrin can be regulated by the redox environment. Collectively, our findings suggest that cysteines in gephyrin modulate synaptic localization and clustering as regulatory redox-switches thereby establishing a link between neuronal and mitochondrial activity.

## INTRODUCTION

Neuronal activity is connected to oxygen consumption and the production of reactive oxygen species (ROS)^1^, that were initially considered as neurotoxic^2^. However recently, redox signaling has emerged as an important pathway for modulating synaptic plasticity. For instance, mitochondria-derived ROS enhance neurotransmission at inhibitory synapses^3^, which was attributed to the recruitment of GABA type A receptors (GABA_A_R) to the synapse, but a molecular mechanism explaining this phenomenon remained elusive. Moreover, it is unclear, how synaptic ROS signaling would be facilitated or sustained in the reductive cytosol. Importantly, maintaining the proper relation of ROS production and clearance is crucial for healthy brain function, and disruptions in ROS homeostasis leading to oxidative stress have been linked to various neuronal diseases^4,5^. Consequently, unraveling the underlying molecular link between ROS production and neuronal activity is of great interest for understanding synaptic plasticity and the onset of pathological processes.

One key component at inhibitory synapses is the bifunctional protein gephyrin. Gephyrin is crucial for the activity of molybdenum cofactor (Moco)-dependent enzymes involved in basic metabolism. The conserved N-terminal G-domain and C-terminal E-domain of gephyrin catalyze the last two steps of Moco biosynthesis^6^. In addition, in the CNS gephyrin serves as the main scaffolding protein at inhibitory synapses, where it clusters glycine receptors (GlyRs) and a subset of GABA_A_Rs to facilitate efficient neurotransmission^7,8^. The synaptic clustering requires the self-assembly of gephyrin through E-domain dimerization and G-domain trimerization^9^. The interaction between receptors and gephyrin takes place between the intracellular domain (ICD) of one receptor subunit and the gephyrin E-domain^9^. The gephyrin C-domain links the catalytic G-and E-domain, providing flexibility as well as sites for post-translational modifications (PTMs) that regulate receptor binding and clustering^10,11^. One example is the proteolytic cleavage of gephyrin C-domain by the calcium-dependent protease calpain^12,13^. This irreversible modification is important for gephyrin homeostasis and links excitatory calcium signaling to the shaping of inhibitory gephyrin scaffolds^12^. Moreover, gephyrin participates in a complex network of protein interactions crucial for signal transmission and neuronal plasticity^14^. Thus, dysfunction of gephyrin has detrimental consequences for both metabolic and neuronal functions, ranging from delayed development to seizures and early childhood death^15–17^.

Recent studies revealed that gephyrin regulation by PTMs, including those targeting cysteines, is essential for synaptic plasticity^18,19^. Palmitoylation at Cys212 and Cys284 was identified to increase synaptic gephyrin clusters, while S-nitrosylation resulted in smaller synaptic clusters. However, not all of gephyrin’s surface cysteines have been assigned a specific function. Given that in the absence of specific functions cysteines get lost during evolution^20^, we investigated a potential link between redox-signaling and surface cysteines going beyond the previously reported nitrosylation and palmitoylation.

Redox modifications on cytosolic proteins are often regulated in space and time. Gephyrin is a cytosolic protein that reversibly associates with the postsynaptic membrane forming postsynaptic clusters/microdomains. Recently, liquid-liquid phase separation (LLPS) has emerged as a novel mechanism for creating subcellular microdomains influencing pre- and post-synaptic functions^21–23^. LLPS has been proposed as a possible mechanism for gephyrin multimerization^22^. In addition, the ability of gephyrin to undergo LLPS may be influenced by redox-changes or may impact gephyrin’s accessibility to redox signals. Initially, it was found that interaction with an artificial multimeric binding partner is necessary to enable gephyrin LLPS ^22^. In the previous study, a dimeric model consisting of two ICDs of the GlyR β-subunit was utilized. In this study, we employed two different receptor models to study the binding and LLPS properties of gephyrin in relation to its redox state. One model is a newly developed pentameric and soluble fusion protein of yeast lumazine synthase (LS) and five ICDs of the GlyR β-subunit^24^, while the other is the established monomeric peptide of the GlyR β-subunit ICD^25^.

Our results demonstrate a dynamic redox-dependent regulatory mechanism for synaptic gephyrin clustering: Oxidation strengthened clustering, a process that was reversed by reduction. As result, the number of receptor binding sites increased, which was consistent with a previously reported redox-dependent enhancement of inhibitory neurotransmission^26^. We showed that LLPS forms the basis for creating oxidative microdomains, with H_2_O_2_ derived from mitochondria being a potential source for gephyrin oxidation. In conclusion, our study establishes a link between inhibitory neuronal and mitochondrial activity and offers a feedforward pathway through redox-dependent synaptic clustering of gephyrin.

## RESULTS

### Surface cysteines of gephyrin are required for synaptic function

A crucial aspect of inhibitory synaptic transmission is the ability of gephyrin to build a scaffold by multimerization. To test whether the oligomeric state of gephyrin is dependent on cysteines and if those are redox-sensitive, we tested whether gephyrin presents redox-sensitive behavior in native tissue. Therefore, we perfused mice with N-ethyl-maleimide(NEM)-containing PBS to stabilize free thiols and avoid unspecific oxidation. In subsequent Western blot analysis, we either used the reductant dithiothreitol (DTT) or omitted it, allowing to compare the presence of redox-dependent gephyrin species (Figure 1A). In all tissues, the absence of a reductant resulted in bands corresponding to a molecular weight of approximately 300 kDa representing multimeric gephyrin species, whereas reduction of gephyrin led exclusively to monomeric gephyrin. The fractional abundance of multimeric gephyrin was approximately 20% (Figure 1B). Based on this finding, we hypothesized that gephyrin cysteines are redox-sensitive forming intermolecular disulfide bridges and thereby influence multimerization. Multimerization is a crucial parameter for the synaptic function of gephyrin^27^. Thus, we wondered which surface cysteines of gephyrin could be important for multimerization.

**Figure 1:**
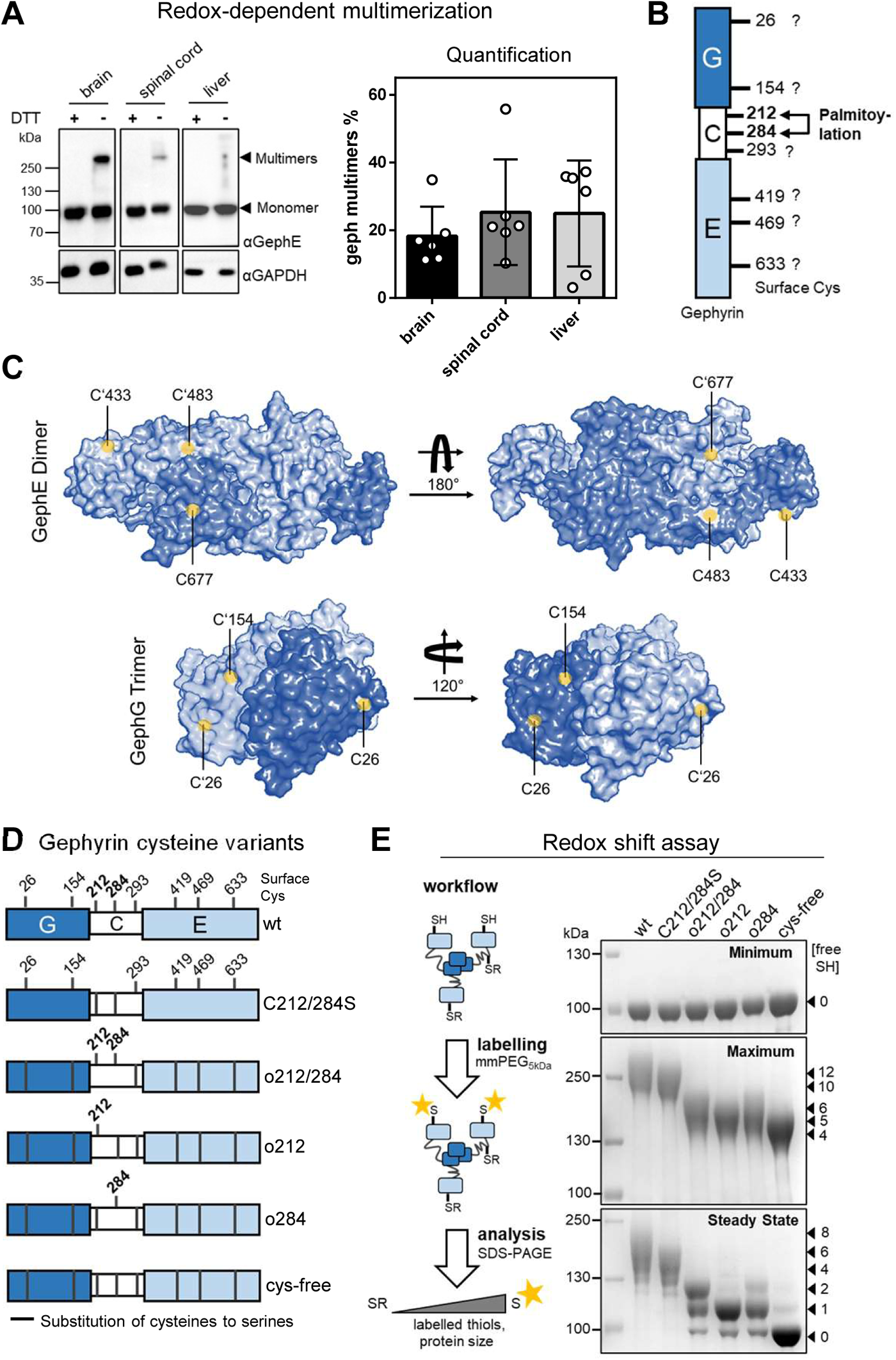
Role of Surface-exposed Cysteines of Gephyrin. A) Western Blot analysis of tissue lysates generated from mice, that have been perfused with NEM. Samples of reduced (DTT) or non-reduced lysates (load of 45 µg). Monomeric and multimeric gephyrin marked with arrows. Quantification of gephyrin multimer bands from A, including six mice samples in total. Abundance in relation (%) to complete gephyrin amount loaded in one sample. B) Domain structure of gephyrin. G=G-domain; C=C-domain; E=E-domain. Positions of the eight potential surface-exposed cysteines highlighted. C) Crystal structures of trimeric G-domain (1IHC^25^) and dimeric E-domain (2FU3^28^). Surface cysteines highlighted in yellow. D) Genetically engineered cysteine variants used in this study. Cysteines were substituted with serines at indicated positions. E) Redox Shift Assay with the cysteine variants. Workflow: In this assay free thiols (-SH) are labelled with a maleimide-PEG-5kDa-tag (mmPEG) increasing the size in SDS-PAGE. Coomassie staining: Amount of labelled, free-thiols cysteines indicated with arrows. As controls mmPEG and SDS were left out (Minimum) and both were added to additionally label the 4 internal cys (Maximum). For the steady state redox-status mmPEG but no SDS was used.

Crystall structures of G- and E-domain^25,28^, as well as the identification of palmitoylation^19^, assigned cysteines at position 26, 154, 212, 284, 419, 469 and 633 as surface-exposed (Figure 1 B and C). Interestingly, those cysteine residues are conserved in orthologues harboring a central nervous system (Figure S1), while other proteins such as Cnx1 in plants or bacterial orthologues lack those cysteines. In addition, Cys293 within the C-domain is considered as surface-exposed in the absence of structural information and a prediction as an intrinsically unordered sequence within the protein^11^. To investigate the potential role of exposed surface cysteines in redox regulation of gephyrin function, we generated cysteine variants by the substitution to serine (Figure 1D). Previously, Cys212 and 284 were identified as targets of palmitoylation^19^. In this study, each of the other eight surface cysteines was individually substituted with serine. These variants were expressed in cultured primary hippocampal neurons, and the synaptic localization of gephyrin was analyzed. Except for the palmitoylated Cys212 and 284, none of the other substitutions resulted in any noticeable effect. This suggested that the other surface cysteines were individually dispensable. To test whether more than one cysteine outside the palmitoylation motif may have a functional role, we generated a gephyrin variant containing only Cys212 and 284 (o212/284) by replacing all other surface cysteines (Cys26, 154, 293, 419, 469, 663) with a serine. Additionally, we generated variants with remaining Cys212 (o212) and Cys284 (o284), respectively, and one variant without any surface-exposed cysteine, which we called “cys-free”. Importantly, in all variants the four internal cysteines (Cys74,133, 635, 676) were unchanged.

The respective gephyrin variants were expressed and purified to homogeneity (Figure S2A). All variants showed similar secondary structure distribution as judged by CD-spectroscopy (Figure S2B). Besides that, we observed similar expression in COS7 cells (Figure S3A and B) and functional enzymatic activity (Figure S3C).

With the purified variants we performed a redox-shift assay (Figure 1E) to determine free thiols by the attachment of a maleimide conjugate with a PEG-tag (mmPEG_5kDa_), which requires an accessible, free thiol. Each labelled cysteine increased the size of gephyrin by approximal 10 kDa. Note that, internal Cys74, 133, 635 and 676 are accessible in the unfolded protein only (maximum shift). Thus, we confirmed that each individual cysteine among the eight predicted ones were indeed surface exposed, regardless of the presence of other cysteines (Figure 1E).

Based on the observed redox-dependent multimerization and redox-shift data, we anticipated a critical role for the eight surface-exposed cysteines in gephyrin. We focused on the scaffolding function of gephyrin in neurons. First, we performed Western blot and RT-PCR analysis demonstrating equal levels of expression for all variants (Figure S4). Afterwards, we expressed EGFP-tagged wt gephyrin, the variant lacking the palmitoylation sites (C212/284S), and our newly generated variants (o212/284 and cys-free) in hippocampal neurons (Figure 2A). Subsequently, we assessed synaptic gephyrin cluster size (Figure 2B). The o212/284 variant did not exhibit a wt-like phenotype. The disturbance of synaptic clustering was similar to that observed in the C212/284S variant, as the synaptic cluster size was significantly reduced. This finding led us to conclude that the other surface cysteines serve an unidentified function, distinct from that of palmitoylation^19^. Furthermore, this unknown function appears to be dependent on multiple cysteines, as single substitutions that were studied before did not show any significant impact on synaptic function^19^. Moreover, the cysteine-free variant was in comparison to wt gephyrin significantly impaired, which was comparable to the o212/284 variant underlying the importance of all surface-exposed cysteines.

**Figure 2:**
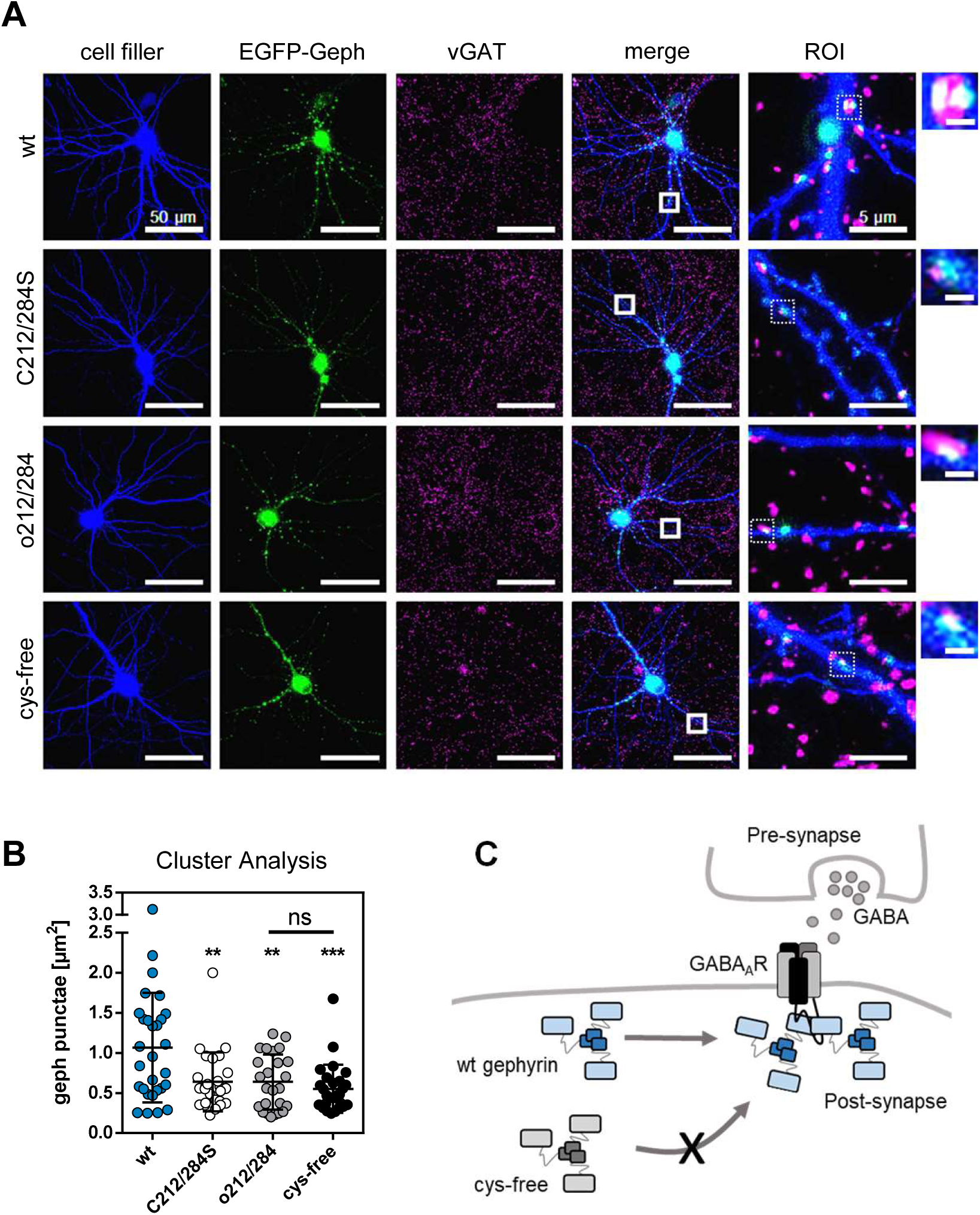
Role of the Surface-Exposed Cysteines for Gephyrin Function. A) Primary, hippocampal neurons transfected with EGFP-gephyrin (green) and tdTomato (cell filler, blue) (scale bar 30 µm). Synaptic termini identified by vGAT staining. Dendrites enlarged (scalebar 5 µm) with increments of respective synaptic gephyrin clusters (scale bar 1 µm). B) Cluster analysis of synaptic gephyrin. Significance tested by 1way ANOVA *F*(3,103)=7.240, *p*=0.0002. *Bonferroni* post-hoc test in comparison to wt: C212/284S: *p*=0.0033 (**); o212/284: *p*=0.0035 (**), cys-free: *p*=0.0002 (***). *Bonferroni* post-hoc test of o212/284 and cys-free *p*=0.9368 (ns). C) Scheme of gephyrin clustering at GABAergic synapses. Cys-free gephyrin shows disturbed synaptic clustering.

In summary, our data demonstrate that at least eight cysteines in gephyrin are surface exposed and play an important role in synaptic clustering (Figure 2C). Furthermore, the underlying mechanism, responsible for the multimerization of gephyrin at synapses relies on the cooperative action of multiple cysteines, which is in line with the identification of redox-dependent multimers found in native tissue.

### Gephyrin undergoes reversible multimerization through disulfide bridge formations

Given that an unidentified mechanism involves the cooperation of cysteines and regulates the density of gephyrin at inhibitory synapses, we proposed the possibility of an oxidation-induced disulfide-bridge formation in gephyrin clusters. To test this hypothesis, we treated recombinantly expressed gephyrin derived from *E. coli* with either the oxidant diamide or the reductant Tris-2-carboxyethyl-phosphine (TCEP) and analyzed the multimeric state of gephyrin using size-exclusion chromatography (SEC). As a negative control, we used cysteine-free gephyrin. Upon oxidation, wt gephyrin displayed a significant increase in high-order multimers, which was not observed in the cysteine-free variant (Figure 3A and B). This led us to conclude that cysteines mediate a mechanism triggering multimerization. To confirm that disulfide bridges are responsible for gephyrin multimerization, we subjected the oxidation-induced multimers to treatment with DTT, which effectively reversed the formation of multimers back to their initial non-oligomeric state prior to oxidation. In addition, we collected samples of the different SEC peaks and prepared them for SDS-PAGE analysis under reducing or non-reducing conditions (Figure S4A and B). We found that reduction and denaturation of all samples including those of the oxidation-induced high-order multimers, resulted in monomeric gephyrin species at around 100 kDa. In contrast, without reductant prior to SDS-PAGE, monomeric gephyrin species disappeared. For the cysteine-free variant, no difference was observed before and after oxidation.

**Figure 3:**
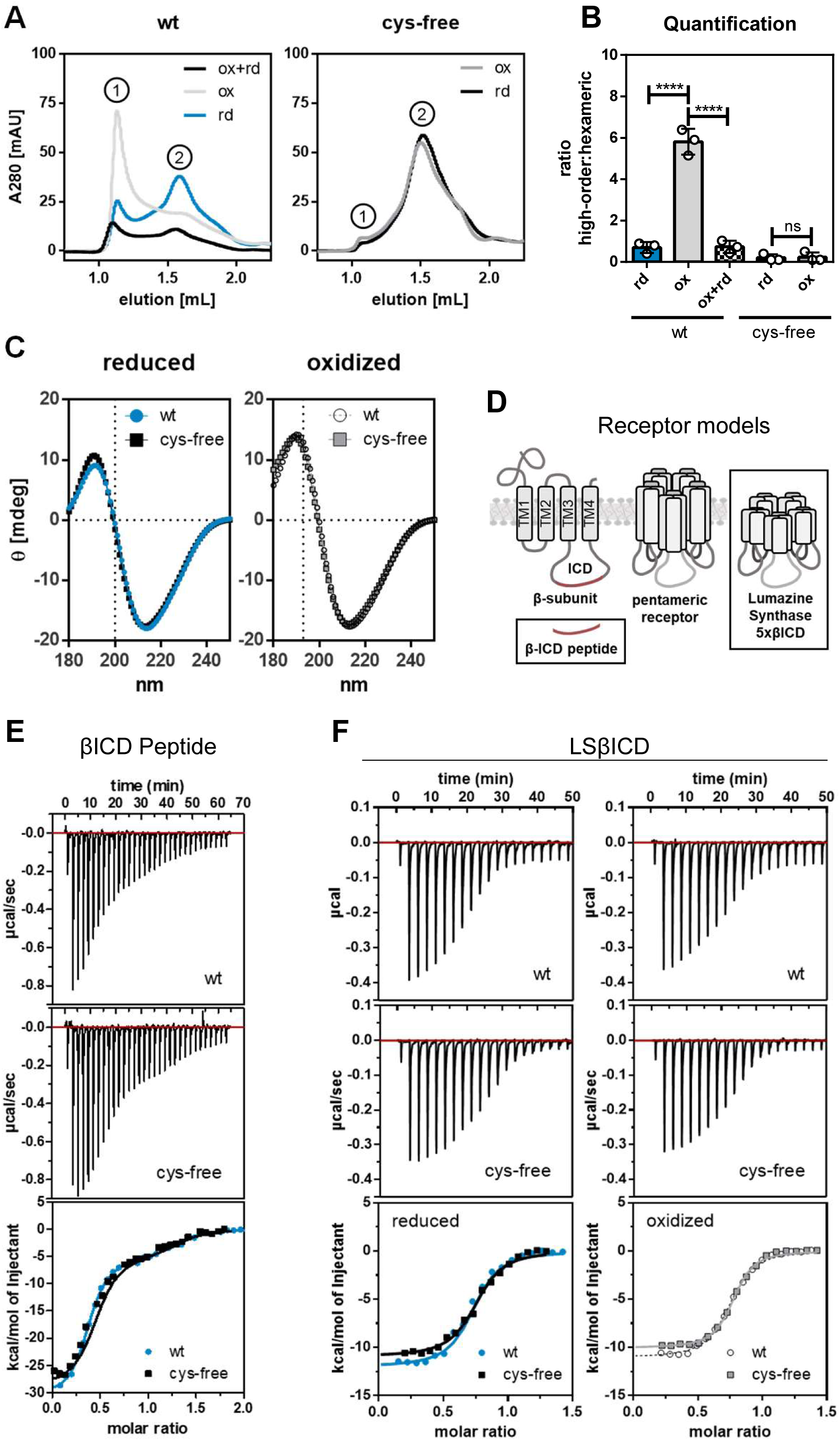
Reversible Redox-Dependent Oligomerization of Gephyrin *in vitro* and *in vivo*. A-B) Size exclusion chromatography (SEC) with oxidized or reduced gephyrin wildtype (wt) or cysteine-free (cys-free) variant. A) Chromatograms with peaks of high-order multimers (1) or hexameric gephyrin (2). Reduced runs in blue (wt) or black (cys-free), oxidized in white (wt) or grey (cys-free) and oxidized peak collected and subsequently re-reduced in black (wt). B) Quantification of the ratio of the peak heights of three different runs. Significance tested by 1way ANOVA *F*(4,10)=137.2, *p<*0.0001, *Bonferroni* post-hoc test: *p<*0.0001 (****); *p*>0.9999 (ns). C) CD Spectroscopy. Spectra of oxidized or reduced gephyrin wt or cys-free variant. Vertical dotted line indicates HT threshold of 800 mV. D) Schematic illustration of receptor models used for binding studies. The glycine receptor (GlyR) is modelled by either a short peptide of the intracellular domains of the GlyRβ-subunit (βICD) or by the fusion of soluble, pentameric lumazine synthase (LS) with five βICDs. E) Isothermal colorimetry (ITC) experiment titrating βICD-peptide into gephyrin wt or cys-free variant. Experiment performed under reductive conditions. F) ITC titrating LSβICD into gephyrin wt or cys-free variant. Experiments were performed under non-reductive conditions with either pre-reduced or pre-oxidized gephyrin.

In conclusion, we show that the mechanism behind redox-dependent multimerization of gephyrin is likely the induction of intermolecular disulfide bridges by oxidation, which can be reversed by reduction.

### Gephyrin oxidation increases liquid-liquid phase separation

We identified that gephyrin multimerization was reversible and inducible by oxidation *in vitro* and we observed redox-dependent multimers *in vivo*. Next, we asked whether these oxidized, multimerized gephyrin species retain their functionality. Given that multimerization is a crucial feature of gephyrin, we sought to investigate the molecular consequences in respect to its biochemical properties, such as structure and receptor binding.

Firstly, we examined the secondary structure and binding capability of both wt gephyrin and the cysteine-free variant under oxidative and reductive conditions. Overall, we observed no alterations in the secondary structure, regardless of whether all cysteines were replaced with serines or if gephyrin underwent reduction or oxidation (Figure 3C). Similarly, we found no changes in the binding properties of gephyrin towards two glycine receptor models βICD and LS βICD^24,25^ using isothermal titration calorimetry (ITC) experiments (Figure 3D-F). For both, wt and cys-free variant, the interaction with the βICD peptide occurred in a two-site binding mechanism (Figure S5C). The stoichiometry observed indicated approximately one gephyrin trimer interacting with one βICD peptide. The dissociation constant K_D_ was in the low micromolar range (wt = 0.041±0.010 μM; cys-free = 0.063±0.018 μM). The binding mechanism of gephyrin wt and cys-free variant with pentameric LS βICD followed a one-site binding model (Figure S5D) yielding a stoichiometry of one LS βICD pentamer to approximately two gephyrin trimers, which was observed for wt and cys-free variant independent of the redox-state. The K_D_ was in the low molecular range as well (wt^rd^ = 0.320±0.042 μM; wt^ox^ = 0.295±0.041 μM; cys-free^rd^ = 0.373±0.041 μM; cys-free^ox^ = 0.291±0.032 μM). Consequently, we concluded that the presence of disulfide bridges did not alter the secondary structure or binding properties of gephyrin, while it changed its oligomeric state.

It would be plausible that oxidation-induced multimerization could lead to tighter packing of gephyrin clusters. Importantly, the quantity of gephyrin present at synapses and the protein density of clusters are crucial factors determining the number of receptor binding sites. Consequently, clustering can impact synaptic plasticity and neurotransmission. In line with this notion, we investigated in which manner oxidative processes would occur and be maintained, since gephyrin is a cytosolic protein and the cytosol represents a reductive environment. One possibility is the ability to form phase-separated droplets in the presence of a multimeric binding partner.

To investigate LLPS of gephyrin in relation to its redox-state, we used the pentameric receptor model LSβICD^24^ (Figure 4A). Following coincubation of a 2:1 molarity of LSβICD to oxidized gephyrin, we found a significantly stronger sedimentation with nearly 100% of all protein in the sedimented fraction, while the majority of reduced gephyrin stayed in the soluble supernatant. In comparison, oxidized and reduced cysteine-free variants remained in the soluble similar to reduced wt gephyrin (Figure 4B). This finding supports the idea that oxidized gephyrin forms high-order oligomeric networks in the presence of oligomeric ligands thus resulting in sedimentation.

**Figure 4:**
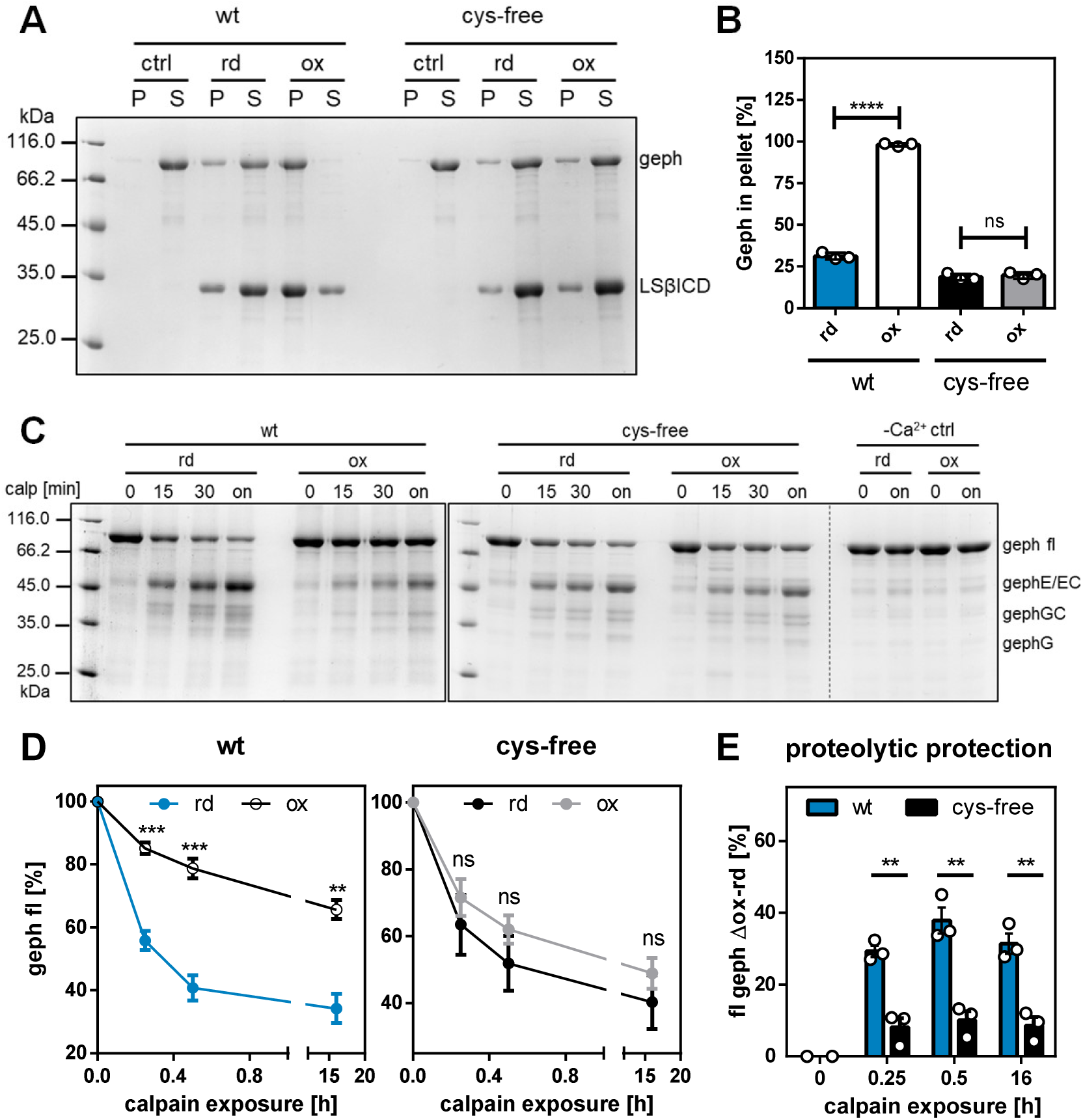
Sedimentation and Calpain Cleavage Assays of oxidized and reduced Gephyrin. Oxidized (ox) or reduced (rd) gephyrin wt and cysteine-free were used for sedimentation (A-and calpain cleavage (C-E) assays. Sum of three replicates. Error bars determined by standard deviation. A-B) Successful interaction with LSβICD causes droplet formation (liquid-liquid phase separation), these droplets are sedimented by centrifugation. Sedimented protein is found in the pellet (P) fraction, non-phase separated protein stays in the supernatant (S). A) Coomassie Staining of fractions. B) Quantification of gephyrin band intensities. Relative abundance (%) normalized to sum of protein in pellet and supernatant of the corresponding fractions. Significance tested by 1way ANOVA *F*_3,8_=1,328, *p*<0.0001. Bonferroni post-hoc test: wt: *p*<0.0001 (****); cys-free: *p*=0.9773 (ns). C-D) Exposure of gephyrin to the protease calpain for different times. C) Coomassie staining. Degradation products of gephyrin indicated at the different heights. fl=full-length; gephE=E-domain; gephEC=E-domain with C-domain fragments; gephG=G-domain, gephGC=G-domain with C-domain fragments. In the control, calcium was absent. D) Quantification of full-length (fl) gephyrin in each sample. Relative abundance (%) normalized to the start point 0. Significance between rd and ox determined by *t-*test and significance level alpha corrected by *Bonferroni* (/3): wt t1:*p=*0.0001 (***); t2:*p*=0.0002 *(***);* t3*:p*=0.006; cys-free: t1*:p*=0.2535 (ns); t2:*p*=0.1292 (ns); t3:*p*=0.1843 (ns). E) Comparison of proteolytic resistance. Relative abundance of fl geph in the reduced samples was subtracted from that of the oxidized ones (Δox-rd). Significance tested by t-test and significance level alpha *Bonferroni* corrected (/3): t1:*p*=0.0022 (**); *p=*0.0031 (**); *p=*0.0033 (**).

Calpain-mediated cleavage is an important mechanism for the regulation of gephyrin clustering at synapses^12^. Besides calcium levels, the access to calpain cleavage sites determines the rate of proteolysis, which could be altered through redox-dependent multimerization. Therefore, the ability of calpain to cleave reduced or oxidized gephyrin was tested *in vitro* (Figure 4C-E). The sensitivity of oxidized gephyrin to calpain cleavage was significantly reduced by approximately two-fold, while there was no significant difference between reduced wt gephyrin and oxidized or reduced cysteine-free gephyrin (Figure 4D). Analyzing the difference in remaining full-length, undigested gephyrin between oxidized and reduced protein (Δox-rd), showed that wt gephyrin exhibited significant proteolytic resistance after oxidation even after short exposure towards calpain (Figure 4E). In contrast, the cysteine-free variant was equally digested as reduced wt gephyrin, regardless of whether it was in the reduced or oxidized state.

In summary, our results show that structural properties of gephyrin that are influenced by its oligomeric state, such phase separation as well as proteolytic sensitivity, were redox-dependent.

### H_2_O_2_ mediates synaptic clustering of gephyrin *in cellulo*

To address the question of how gephyrin wt in comparison to the cys-free variant could be oxidized to multimerize via disulfide-bridge formation in neurons, we investigated H_2_O_2_, which is a commonly known oxidant emerging from different sources in the cell. Therefore, we expressed a cytosolic version of a D-amino acid oxidase (DAAO) with an mScarlet-tag in hippocampal neurons and tested the effect on endogenous gephyrin clusters (Figure 5A) and afterwards on recombinant cys-free and wt gephyrin (Figure 5B). DAAO enzyme produces H_2_O_2_ by the conversion of D-amino acids to the corresponding α-keto acids (Figure 5A), thus we treated our cells with D-Met. As a negative control we generated a less active variant of DAAO by substituting Arg285 to Lysin^29^. Following cell treatment, we stained for the presynaptic marker vGAT and analyzed its co-localization with endogenous gephyrin to identify synaptic gephyrin species.

**Figure 5:**
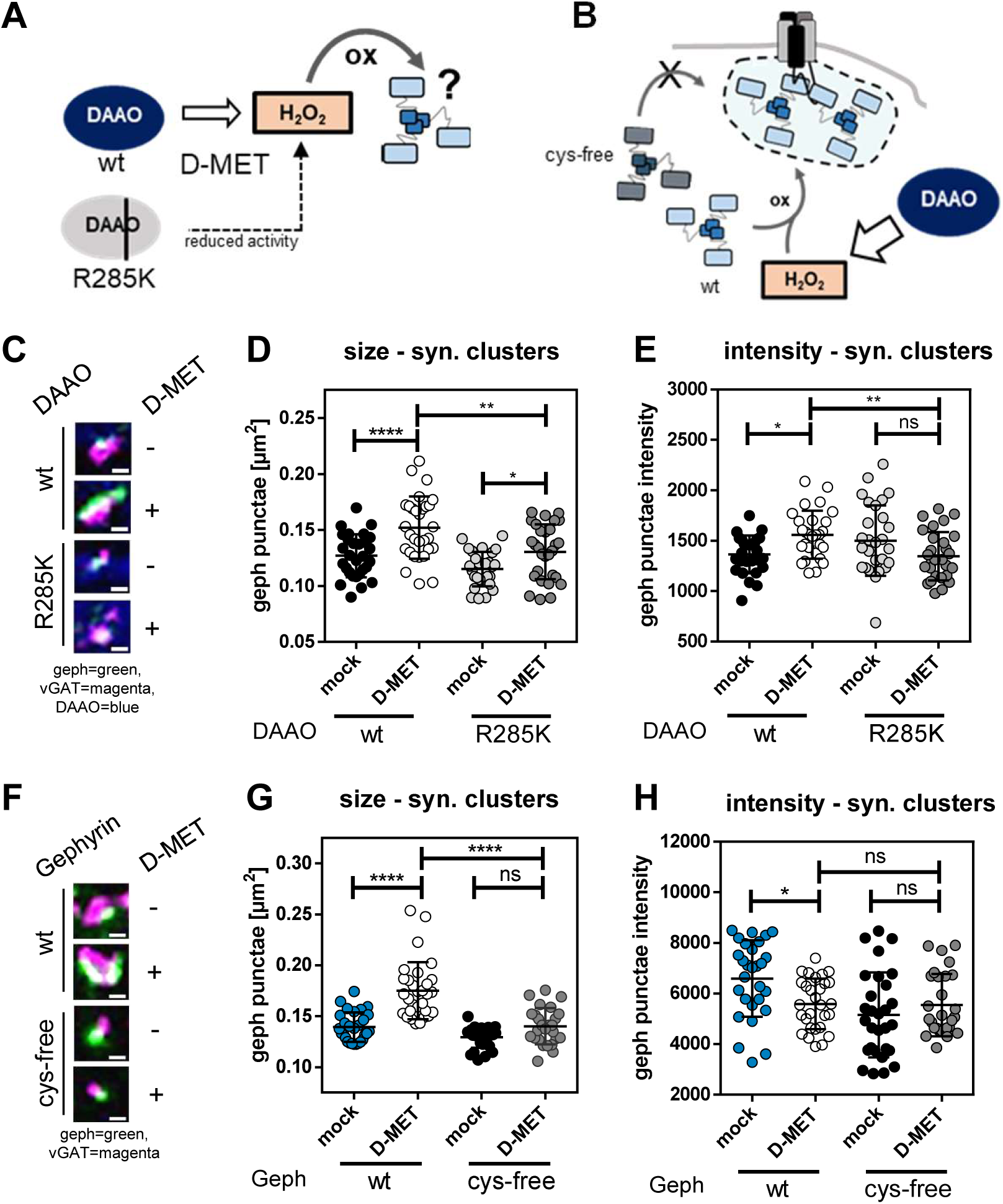
H_2_O_2_ drives Gephyrin Oligomerization at Synapses. Expression of cytosolic mSclt-DAAO and mEGFP-Geph in primary, hippocampal neurons. Treatment at DIV14 with 5 mM D-MET. Analysis of cluster sizes [µm^2^], intensity [au] and co-localization with vGAT (=synaptic clusters) of gephyrin. A) Schematic description of H_2_O_2_ production of DAAO wt upon D-methionine acid treatment. DAAO mutant R285K is inactive. Can H_2_O_2_ oxidize gephyrin and influence its oligomerization? B) Schematic description of influence of DAAO-generated H_2_O_2_ on gephyrin clustering. Is cys-free gephyrin not influenced? C-D) mSclt-DAAO wt and R285K. Analysis of endogenous gephyrin clusters. C) Representative gephyrin clusters (Gephyrin: green, vGAT: magenta, DAAO: blue) (scale bar 0.5 µm). Synaptic termini identified by vGAT co-localization. D) Sizes of synaptic gephyrin. Significance tested by 1way ANOVA *F*(3,113)=14.18, *p<*0.0001. *Bonferroni* post-hoc test: wt vs. wt+D-MET: *p*<0.0001 (****); R285K vs R285K+D-MET: *p*=0.0277 (*), wt+D-MET vs R285K+D-MET: *p*=0.0010 (***). E) Intensity of synaptic gephyrin. Significance tested by 1way ANOVA *F*(3,109)=4.431, *p=*0.0056. *Bonferroni* post-hoc test: wt vs. wt+D-MET: *p*=0.0204 (*); R285K vs R285K+D-MET: *p*=0.0808 (ns), wt+D-MET vs R285K+D-MET: *p*=0.0083 (**). F-H) mSclt-DAAO wt and mEGFP-Geph wt and cys-free variants. Analysis of recombinant mEGFP-gephyrin clusters. F) Representative gephyrin clusters (Gephyrin: green, vGAT: magenta) (scale bar 0.5 µm). Synaptic termini identified by vGAT co-localization. G) Sizes of synaptic gephyrin. Significance tested by 1way ANOVA *F*(3,107)=31.66, *p*<0.0001. *Bonferroni* post-hoc test: wt vs. wt+D-MET: *p*<0.0001 (****); cys-free vs cys-free+D-MET: *p*=0.1252 (ns), wt+D-MET vs cys-free+D-MET: *p<*0.0001 (****). H) Intensity of synaptic gephyrin. Significance tested by 1way ANOVA *F*(3,110)=5.608, *p*=0.0013. *Bonferroni* post-hoc test: wt vs. wt+D-MET: *p*=0.0212 (*); cys-free vs cys-free+D-MET: *p*=0.8567 (ns), wt+D-MET vs cys-free+D-MET: *p*>0.9999 (ns).

Firstly, the ubiquitous and cytosolic localization of DAAO could be demonstrated (Figure 5C, full images in Figure S6A). D-Met treatment caused a significant increase of synaptic cluster sizes by approximately 20% (from 0.1269 to 0.1521 µm^2^) (Figure 5C and D). Additionally, the intensity of synaptic gephyrin was significantly increased by about 14% (from 1 364 to 1 559 a.u.) (Figure 5E). The expression of DAAO R285K variant in combination with D-Met treatment had a significant increase on gephyrin cluster size as well, but the effect size was smaller (Confidence Interval [-0.011;-0.039]^wt^, [-0.001;-0.030]^R285K^). The effect could be explained by the residual activity of the R285K mutant^29^. However, no effect of the treatment in combination with the R285K mutant on the intensity of gephyrin was observed. Based on this observation, we conclude that the effects on gephyrin resulted mainly from the enzymatic activity of DAAO, the production of H_2_O_2_. Since increase of cluster size and intensity are signs of gephyrin recruitment to the synapse, we concluded that H_2_O_2_ is able to trigger synaptic multimerization of gephyrin *in cellulo*.

Next, we verified that H_2_O_2_ has a direct effect on the cysteines of gephyrin to support the hypothesis of gephyrin oxidation-mediated multimerization via disulfide-bridges (Figure 5B). Thus, we combined the expression of DAAO and subsequent D-Met treatment with the overexpression of mEGFP-tagged gephyrin (Figure 5F, full images in Figure S6B). As expected, overexpressed wt gephyrin showed a significant increase of about 25% (from 0.1393 to 0.1750 µm^2^) in cluster size following D-Met treatment, while cys-free gephyrin showed no effect (Figure 5 F and G). Strikingly, the intensity of synaptic mEGFP-gephyrin wt upon DAAO-mediated H_2_O_2_ production was decreased by -15% (from 6 591 to 5 588 a.u.), but that of cys-free gephyrin was unchanged (Figure 5H). In this set up, oxidation extended existing gephyrin clusters in size but did not increase the density within clusters.

In conclusion, we could show that DAAO-mediated H_2_O_2_ promotes synaptic gephyrin cluster formation. The mechanism involves surface cysteines of gephyrin since cysteine-free gephyrin did not react to cellular H_2_O_2_ formation.

### Mitochondria-derived ROS are a source of gephyrin oxidation

Lastly, we were looking for a more natural source of oxidation since the expression of ubiquitous DAAO is an artificial approach. Such a natural source could be ROS generated by mitochondria (mROS). Although there are more pathways generating ROS, we chose mitochondria, since neurons are mitochondria-rich cells requiring ATP and thus active mitochondria^30–32^. In turn, mitochondria with oxidative phosphorylation activity, likely produce ROS such as superoxide radicals, that are quickly turned over to H_2_O ^33^.

ROS production was triggered by the treatment of neurons with antimycin A. Antimycin A inhibits complex III of the respiratory chain in mitochondria leading to the accumulation of electrons^34^. These electrons react with oxygen to ROS^35^. Additionally, we introduced DTT treatment to reverse mROS-dependent oxidative stress (Figure 6A). Again, we monitored first endogenous gephyrin and the presynaptic marker vGAT (Figure 6C, full images in Figure S7A). Comparable to the results using DAAO as an oxidation source, mROS production led to a significant increase of gephyrin average cluster size by about 13% (from 0.1808 to 0.2051 μm^2^) and significantly increased gephyrin intensity by about 25% (from 810.7 to 1 020 a.u.) at synapses (Figure 6D and E). This effect was reversible following DTT treatment (to 0.1851 μm^2^ and 890.1 a.u.).

**Figure 6:**
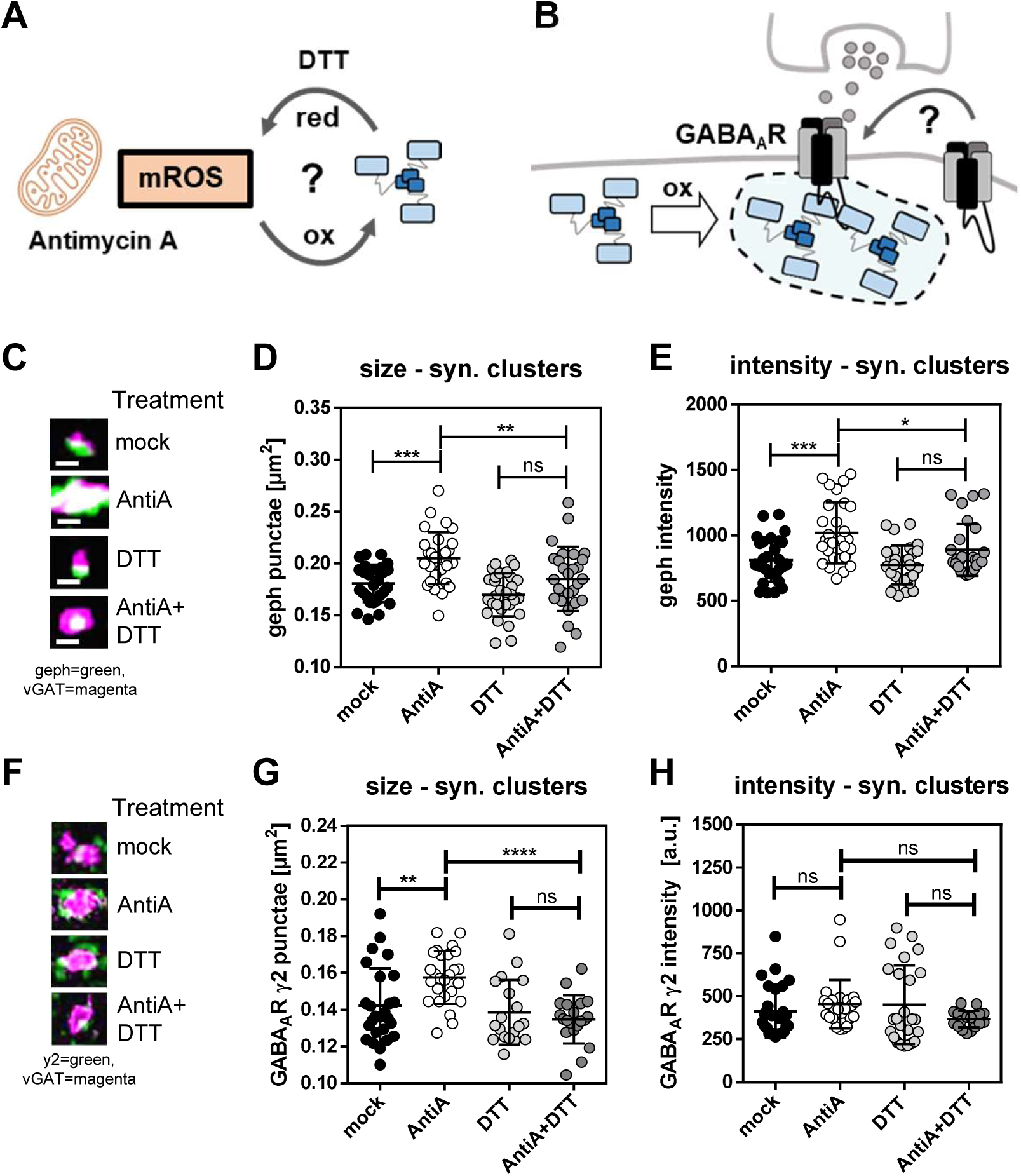
mROS reversibly drive Gephyrin Oligomerization and the Recruitment of GABA_A_R γ2 Subunits. Treatment of primary hippocampal neurons at DIV14 with 1 μM Antimycin A (AntiA) and/or 5 μM DTT. Analysis of gephyrin or GABA_A_R γ2 subunit cluster sizes [µm^2^], intensity [au] and co-localization with vGAT (=synaptic clusters). A) Schematic description of mitochondrial ROS (mROS) production through Antimycin A treatment. Do mROS oxidize gephyrin and can DTT reverse this by reduction? B) Schematic description of influence of H_2_O_2_ on GABA_A_R (containing γ2 subunits) clustering. Does gephyrin oxidation lead to more synaptic receptors? C-E) Analysis of endogenous gephyrin. C) Representative gephyrin clusters (Gephyrin: green, vGAT: magenta, DAAO: blue) (scale bar 0.5 µm). Synaptic termini identified by vGAT co-localization. D) Sizes of synaptic gephyrin. Significance tested by 1way ANOVA *F*(3,112)=11.08, *p<*0.0001. *Bonferroni* post-hoc test: mock vs. AntiA: *p*=0.0006 (***); DTT vs DTT+AntiA: *p*=0.0540 (ns), AntiA vs AntiA+DTT: *p*=0.0065 (**). E) Intensity of synaptic gephyrin. Significance tested by 1way ANOVA *F*(3,111)=9.734, *p<*0.0001. *Bonferroni* post-hoc test: mock vs. AntiA: *p*=0.0001 (***); DTT vs. DTT+AntiA: *p*=0.0676 (ns), AntiA vs. DTT+AntiA: *p*=0.0341 (*). F-H) Analysis of endogenous GABA_A_R γ2 subunit clusters. F) Representative γ2 clusters (γ2 : green, vGAT: magenta) (scale bar 0.5 µm). Synaptic termini identified by vGAT co-localization. G) Sizes of synaptic γ2 clusters. Significance tested by 1way ANOVA *F*(3,87)=8.567, *p<*0.0001. *Bonferroni* post-hoc test: mock vs. AntiA: *p*=0.0037 (**); DTT vs. DTT+AntiA: *p>*0.9999 (ns), AntiA vs. DTT+AntiA: *p<*0.0001 (****). H) Intensity of synaptic γ2 clusters. Significance tested by 1way ANOVA *F*(3,93)=1.432, *p*=0.2384.

Previously, it was reported that antimycin A treatment enhanced inhibitory neurotransmission by the recruitment of synaptically active GABA_A_ R containing γ2 and α3 subunits to the synapse^26^. Oxidation-mediated multimerization of gephyrin potentially leads to more available receptor binding sites, which could be an explanation for the recruitment of GABA_A_ Rs (Figure 6B). To test this functional link, we again treated hippocampal neurons with antimycin A to produce mROS and stained for synaptic GABA_A_ R γ2 subunits (synaptic = co-localization with vGAT) (Figure 6F-H, full images in Figure S7B). Upon antimycin A treatment, the average sizes of synaptic GABA_A_ R γ2 punctae significantly increased by about 11% (from 0.1421 to 0.1576) (Figure 6G). This effect was again reversible following DTT treatment. The intensity of γ2 punctae was unchanged in all conditions (Figure 6H).

Taken together, mROS production enhanced synaptic gephyrin clustering, which was reversible by reduction. This finding indicates that the underlying mechanism of multimerization is redox-dependent and reversible. Since cysteine-free gephyrin did not respond to mROS, we propose that oxidation-driven multimerization is mediated via disulfide-bridge formation of gephyrin surface cysteines. The multimerization of gephyrin at the synapse is important for the supply of receptor binding sites. We could show that mROS-mediated oxidation of gephyrin strengthens the recruitment of synaptically active GABA_A_ Rs containing γ2 subunits. This likely finally supports inhibitory neurotransmission thus linking mitochondrial to neuronal activity through redox-dependent multimerization of gephyrin.

## DISCUSSION

Our study provides new insight into the mechanism of gephyrin-dependent inhibitory synapse formation. First, we identified surface exposed cysteines *in vitro* and demonstrated their sensitivity to redox changes. As a result, we found that the multimerization of gephyrin was regulated by oxidation and reduction. We identified redox-dependent multimers *in vivo* and induced synaptic multimerization *in cellulo* through DAAO-mediated H_2_O_2_ and antimycin A treatment targeting mitochondria. Oligomerization is a crucial aspect of gephyrin scaffolds, significantly impacting the plasticity of inhibitory synapses^27,36^. Consequently, the oligomeric state influences the number of available receptor binding sites and the sensitivity of gephyrin to proteolytic cleavage^37^. Changing the oligomeric state by oxidation enhanced clustering, while reduction reversed this process (Figure 7). In aggregate, this study provides a novel role of multiple surface-exposed cysteines in gephyrin, which were found to be conserved in orthologues with a central nervous system while other orthologs in plants, yeast, fungi and prokaryotes lack those residues (Figure S4).

**Figure 7:**
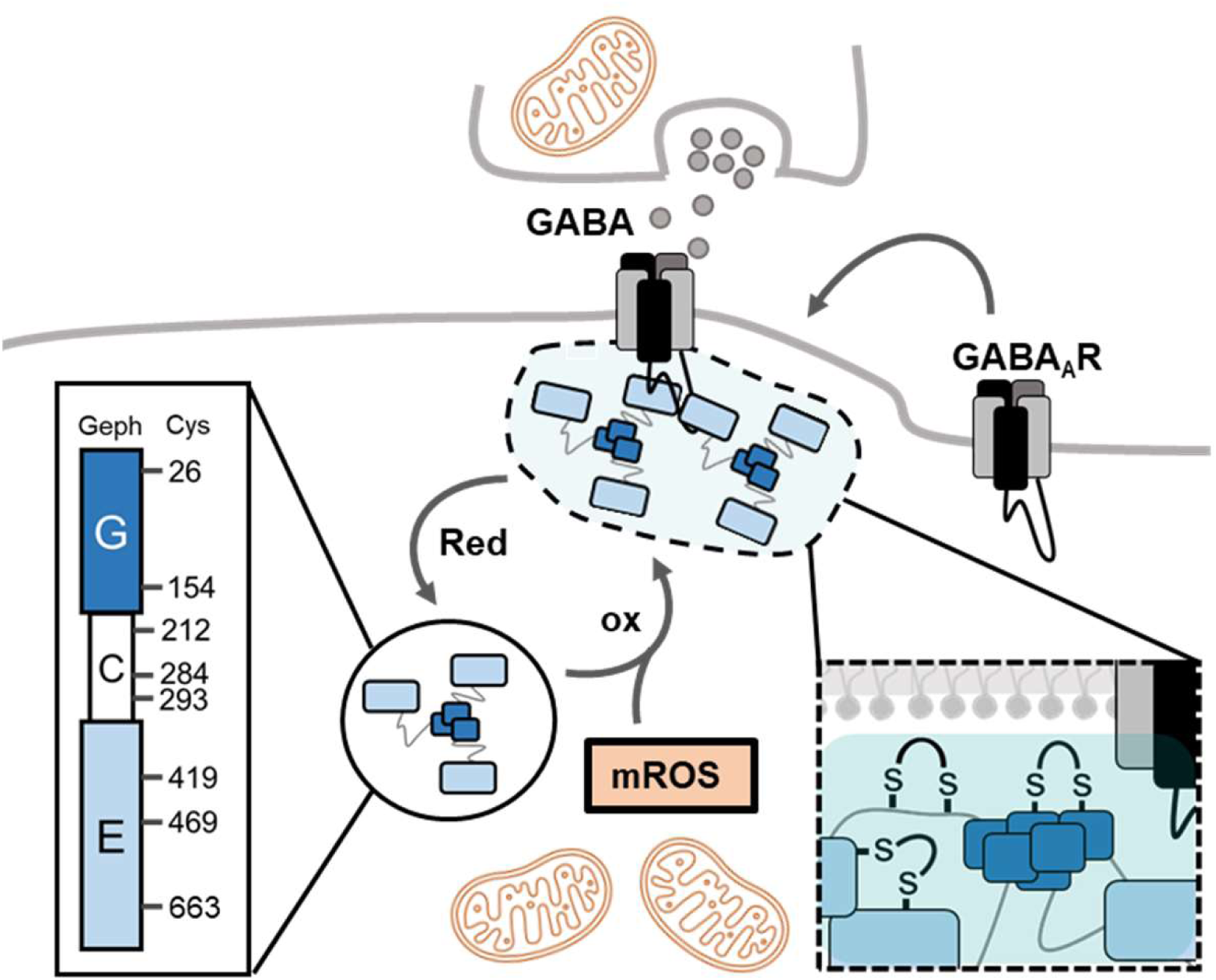
Redox-Regulation of Synaptic Gephyrin Clustering. Hypothesized model of redox-dependent gephyrin oligomerization at inhibitory GABAergic post-synapses. Gephyrin harbors eight surface-exposed cysteines that are redox-sensitive. mROS-mediated oxidation leads to the formation of disulfide bridges (Enlargement highlighted with box surrounded by dotted lines) linking gephyrin molecules increasing gephyrin oligomerization. This oxidation is reversible by reduction and allows for dynamic shuttling of gephyrin localization. LLPS (highlighted in light-blue) might offer an oxidative microdomain facilitating disulfide bridge formation. Ultimately, gephyrin oligomerization at synapses offers binding sites for GABA_A_Rs strengthening inhibitory signal transmission.

Our data suggests that the molecular basis for redox-dependent multimerization of gephyrin is the formation of disulfide bridges. In comparison to PSD95, the excitatory pendant, multimerization by the formation of disulfide bridges was also detected *in cellulo* and *in vivo*^38^. Recently, the disruption of a cysteine linkage in PSD95 has been linked to an epileptic phenotype in rats^39^ highlighting the importance of redox-regulation of neuronal scaffold proteins.

Cysteine oxidation in gephyrin could be reversed by reduction, which is essential for enabling dynamic changes and maintaining plasticity. Importantly, cysteine oxidation did not influence the secondary structure of gephyrin and its binding to the receptor, but it altered the density of gephyrin at the synapse, manipulating the availability of receptor binding sites. Since the y2 subunit is considered crucial for synaptic GABA_A_Rs^7^, we provide an explanation for the increase in inhibitory neurotransmission caused by antimycin A in previous studies^26^: redox-dependent multimerization of gephyrin, which recruited GABA_A_R-containing y2 subunits.

Although, redox-dependent regulation of synaptic proteins has previously been investigated, important questions remain open such as the source of oxidation. Given that neurons are rich in mitochondria and mitochondria are located in close proximity to the synapse^31^, we propose that mitochondria-derived ROS could serve as a source for oxidation. In this sense, gephyrin could provide the functional link between mitochondrial and neuronal activity. Highly active neurons require ATP production mainly derived from mitochondrial respiration being accompanied by the formation of mROS, which is rapidly converted to H_2_O_2_ ^40–42^. H_2_O_2_ can reach the synapse and trigger gephyrin recruitment and network expansion, thereby recruiting additional receptors to further increase neurotransmission. Mitochondria are located in both pre-synaptic and post-synaptic regions, so it is conceivable that mROS can regulate the post-synapse from various angles and in reverse it can impact both sites of the synapse.

Besides mitochondria, there are additional origins of H_2_O_2_, such as the NADPH-oxidase 2 (NOX2) pathway^43^. NOX2 is highly abundant in cells throughout the CNS^44^ and produces superoxide radicals in the extracellular environment, which are subsequently converted to H_2_O_2_. This H_2_O_2_ can re-enter the cell through aquaporins. In a recent study, the effects of this pathway on inhibitory neurotransmission were investigated^45^. It was shown that communication of excitatory and inhibitory neurotransmission is mediated by ROS and nitric oxide, leading to the recruitment of GABA_A_R-containing γ2 and α3 subunits at inhibitory synapses. In contrast to our study, they identified the GABA_A_R-associated protein(GABARAP) as the primary mediator of receptor recruitment, a protein that has been found to interact with gephyrin^46^. Although, it is unclear, whether different ROS-mediated signaling mechanisms can co-exist, it highlights the importance of these mechanisms in regulating neuronal plasticity.

Another open question concerns the maintenance of oxidation processes in the reductive cytosol. In recent studies, LLPS has gained attention as a new principle of subcellular organization^21,47,48^. For gephyrin it was recently demonstrated that it forms liquid droplets in the presence of a multimeric receptor model^22^. Importantly, our data showed that oxidation of gephyrin strengthened LLPS. Assuming that gephyrin could form liquid droplets at synapses within the cell, these microdomains and their reactive cysteines may be shielded from the reductive cytosol allowing the maintenance of their disulfide bridges. Testing the hypothesis that gephyrin is shielded from the reducing cytosol to maintain its oxidized state is challenging *in cellulo* or *in vivo*. Moreover, it remains unclear whether oxidation leads to phase separation or *vice versa*. We showed that the multimerization of gephyrin can be controlled by oxidation and reduction, indicating a reversible and dynamic mechanism. Thus, it is possible that gephyrin could shuttle between phase-separated microdomains depending on its redox state. Additionally, these oxidizing microdomains are likely not a dead end, as reduction was able to reverse oxidation-mediated clustering of gephyrin *in cellulo*. This reversibility is essential to prevent harmful aggregation and facilitate dynamic plasticity.

More and more studies target a possible relation between redox state of cysteines and the ability to undergo LLPS. One example is synapsin 1, which harbors two cysteines directly affecting LLPS in a redox-dependent manner, which is also relevant in aging^23^. A study, that is mechanistically comparable to our work, was reported for oleosin, where the disulfide-bridge formation of artificially added cysteine residues on the protein surface triggered LLPS, while reduction was able to reverse this effect^49^. In general, the concept of a reversible formation of phase-separated microdomains holds great potential and may be a key to understand redox-dependent processes in different cellular environments.

Oligomerization as a function of the redox state of cysteines (including nitrosylation, persulfidation, sulfenylation and disulfide-formation) has been investigated previously^23,40,50^. Such modifications may lead to functional multimers or harmful aggregates depending on the site and degree of oxidation thus reflecting the dose of ROS^51^. Imbalance in redox signaling is linked to aging and neurodegenerative diseases^52^. Our study identifies gephyrin as mediator between mitochondrial activity and inhibitory synapse formation using ROS as signaling mechanism.

## MATERIALS AND METHODS

### DNA/expression constructs

Gephyrin P1 and cysteine variants in pQE80L (Qiagen) were cloned by Gibson assembly using EcoRI into pAAV_hSyn_mScarlet (a gift from Karl Deisseroth, Addgene plasmid #131001) and using EcoRI into pAAV_hSyn_mEGFP (generated from moxBFP, a gift from Erik Snapp, Addgene plasmid #68064; http://n2t.net/addgene:68064; RRID: Addgene_68064). Gephyrin was cloned using PCR and XhoI and KpnI into pEGFP-C2 (Clonetech). The template for the generation of rat Gephyrin P1 constructs was in pQE80L ^53^. The cysteine to serine substitutions were generated by site-directed mutagenesis. DAAO was cloned into pAAV_hSyn_mScarlet through PCR and Gibson Assembly using EcoRV and the R285K mutation was introduced by site-directed mutagenesis. The plasmids pAdDeltaF6 (a gift from James M. Wilson, Addgene plasmid #112867; http://n2t.net/addgene:112867; RRID: Addgene_112867) and pAAV2/1 (a gift from James M. Wilson, Addgene plasmid #112862; http://n2t.net/addgene:112862; RRID: Addgene_112862) were used for virus production. The expression constructs of LSβICD in pQE70 and GlyRβICD in ptYB were described previously ^24,54^.

### Recombinant expression and purification of gephyrin

*E. coli* BL21 Rosetta star was used for the expression of His-gephyrin. Cultures were grown at 37°C at 110 rpm and the expression induced at an OD_600_ of 0.3 with 100 nM IPTG. After expression for 22 h at 18°C and 110 rpm, cells were harvested. To achieve reduced cysteines (free thiols), all solutions were supplemented with reductant until final storage of isolated proteins. The cell pellet was resuspended in lysis buffer (300 mM NaCl, 50 mM Tris/HCl, 5 mM beta-mercapto ethanol, 0.05% Tween20, 1 µg/mL lysozyme, 1x PI (*Roche*), pH 7.4) and stored at -80°C. Pellets were thawed in a water bath and all steps performed at 4°C. Cells were lysed through alternating ultrasound (3 min at 4°C, 30 sec pulse, 30 sec pulse, 40% amplitude) and pressure lysis (EmulsiFlex, 1 000-1 500 bar, 4°C) twice. The homogenate was centrifuged (45 min, 16 000 rpm, 4°C, JLA 16.250 rotor, Beckman-Coulter Centrifuge Avanti J25-01) and the supernatant subjected to Ni-NTA affinity purification (HIS-Select Nickel Affinity Gel (Sigma Aldrich)). All buffers for washing and elution were similar to the lysis buffer, contained no lysozyme, but contained 20 mM, 40 mM and 300 mM Imidazole, respectively. The eluate was further subjected to preparative size-exclusion chromatography (Superdex 200, 120 mL, *GE Healthcare*) using SEC buffer (300 mM NaCl, 50 mM Tris/HCl, 5 mM beta-mercapto ethanol, 0.05% Tween20, 5% glycerol, pH 7.4). The fractions of the peaks eluting around 60-70 mL (Trimers) and 45-60 mL (higher multimers) were collected and concentrated by an *Amicon* filter with a Cut-off of 50 kDa. To remove reductants, the concentrated protein was buffer exchanged by gel filtration (PD10 column) to storage buffer (300 mM NaCl, 50 mM Tris/HCl, pH 7.4) and stored at -80°C.

### Purification of receptor models LSβICD and βICD peptide from *E. coli*

*E. coli* BL21 Rosetta star was used for the expression of His-LSβICD. The expression was performed as previously described^24^, but instead of preparative size exclusion chromatography, the buffer was exchanged by gel filtration using PD10 columns. The final storage buffer was 300 mM NaCl, 50 mM Tris/HCl, pH 7.4.

For the βICD peptide we used *E. coli* ER2566. Cultures grew at 37°C and 110 rpm until the OD600 reached 0.3. Induction was induced by 250 µM IPTG and expression performed for 20 h at 25°C and 110 rpm. Cells were harvested and stored at -80°C in ICD lysis buffer (300 mM NaCl, 50 mM Tris/HCl, 1 mM EDTA, 1x Protease Inhibitor (*Roche*), pH 7.4). Lysis of the thawed cells was performed by alternation of ultrasound (3 min at 4°C, 30 sec pulse, 30 sec pulse, 40% amplitude) and pressure lysis (*EmulsiFlex*, 1 000-1 500 bar, 4°C) twice. After centrifugation for 1 h at 16 000 x g (JLA 16.25 rotor) at 4°C, the supernatant was applied on a pre-equilibrated chitin matrix (*New England Biolabs*) and incubated 1.5 h at 4°C under shaking. The column was washed with 15 CV ICD buffer before 2.5 CV cleavage buffer (300 mM NaCl, 50 mM Tris/HCl, 1 mM EDTA, 50 mM DTT, pH 7.5) was applied an incubated for minimum 24 h at room temperature (rt). The eluate was collected and another 2.5 CV cleavage buffer used for a second elution. The combined eluate was concentrated and filtered by *Amicon* filters with a cut-off of 10 kDa and 3 kDa. The buffer was exchanged to ITC buffer (300 mM NaCl, 50 mM Tris/HCl, pH 7.4) by dialysis over night at 4°C. The purified protein was stored at -80°C.

### Redox shift assay

Typically, 10 μg protein were prepared in a total volume of 20 μL RSA buffer (50 mM Tris/HCl, 5 mM EDTA, pH 7.5) or denaturing RSA buffer (50 mM Tris/HCl, 5 mM EDTA, 2% SDS pH 7.5) and supplemented with 1 mM mmPEG5kDa or buffer as minimum controls. After incubation for 1 h in the dark at rt, samples were used for SDS PAGE. The expected shift size per labeled thiol is about 10 kDa. In denaturing conditions, internal cysteines are exposed to PEGylation additionally.

### *In vitro* redox-modification

Purified proteins were oxidized by incubation over night at 4°C with 1 mM Diamide. Reduction was performed with 1 mM TCEP or DTT. In case the reagent would disturb in further assays, the chemical was removed by gel filtration using PD10 columns.

### Co-sedimentation assay

Recombinant protein was pre-cleared at 17 000 x g for 3 min at rt. We mixed gephyrin and LSβICD 5:10 μM in a final volume of 50 μL of assay buffer (150 mM NaCl and 25 mM Tris/HCl pH 7.5). After incubation of 10 min at rt, samples were centrifuged 10 min at 4°C and 17 000 x g. The supernatant was added in a new tube, supplemented with 15 μL 5x SDS sample buffer. The pellet was supplemented with 50 μL assay buffer and 15 μL 5x SDS sample buffer. Before performing SDS PAGE and Coomassie staining, samples were incubated 5 min at 95°C.

### ITC

Isothermal titration Calorimetry (ITC) experiments were performed by using MicroCal Auto-ITC200 (*Malvern*) at 25°C in ITC buffer (300 mM NaCl, 50 mM Tris/HCl, pH 7.4). We used an injection volume of 1-2 μL, a spacing of 150 sec between injections and an initial delay of 60 sec. The reference power was 5 μCal/sec and the stirring speed 1 000 rpm. The concentrations of LSβICD and GlyRβICD in the syringe were about 200-215 μM and 300-310 μM, respectively and the concentration of gephyrin in the sample cell was 31–35 μM. Parameters and curves were fitted and calculated by the software MicroCal Analysis and Origin7.

### Size exclusion chromatography

Molecular masses of purified gephyrin and variants were detected by analytical size exclusion chromatography (SEC) using a Superose 6 increase 5/10 (*GE Healthcare*). Typically, 4 nmol pre-cleared protein in 50 μL total sample volume were applied. Proteins were separated at 4°C and a flow rate of 0.1 mL/min in SEC buffer (50 mM Tris/HCl, 300 mM NaCl pH 7.4). If desired, fractions of 100 µL were collected and those of corresponding peaks combined and concentrated using *Amicon* filters with a 50 kDa cut-off. The combined protein was then further subjected to redox-modifications, cleared by centrifugation and re-applied on the column.

### Calpain cleavage assay

A mastermix of 50 μL contained 850 pmol redox-modified protein, 10 mM Ca^2+^Cl_2_, 1/2 U/mL calpain. The volume was adjusted using calpain buffer (300 mM NaCl, 50 mM Tris/HCl, pH 7.4). As a control, a calcium-free control was included because calpain is a calcium-dependent protease. At timepoints of 0, 5, 15, 30 min and overnight, 15 µL samples were taken out and the digestion stopped by addition of SDS sample buffer and incubation for 5 min at 96°C. Cleavage products were afterwards analyzed by SDS-PAGE.

### Circular dichroism spectroscopy

Determination of the secondary structure was performed by circular dichroism spectroscopy (CD spec). Typically, 0.3 mg/mL pre-cleared protein in CD buffer (10 mM K_2_HPO_4_/KH_2_PO_4_, pH 7.5) was used and Far-UV spectra recorded with J-715 CD spectropolarimeter (*Jasco*) at 20°C in the range of 180-250 nm using quartz cuvette with 0.1 cm path length. The spectra were processed by the software Spectra Analysis version 1.53.07 of Spectra Manage version 1.54.0. We subtracted the buffer baseline and smoothed three-times by the Savitzky-Golay algorithm with a convolution width of 21.

### Virus preparation

Recombinant virus particles (AAV2/1) were produced in HEK293 cells and isolated by PEG/NaCl precipitation followed by chloroform extraction according to the previously established protocol^55^. The titers were determined by a Gel green® (*Biotium*) protocol^56^ and the fluorescence was detected (507 ± 5 nm excitation and 528 ± 5 nm emission) using a plate reader equipped with monochromators (Tecan Spark).

### Culture of hippocampal neurons and expression of recombinant gephyrin

At E17.5, dissociated primary hippocampal cultures were prepared from C57BL/6NRj embryos. 90 000 cells were seeded on Poly-L-lysine coated 13-mm cover slips in 24-well plates. Neurons grew in neurobasal medium supplemented with B-27, N-2, and L-glutamine (*Thermo Fisher Scientific*). At 8 days *in vitro* (DIV), cells were transfected with 0.4 µg plasmid DNA per construct using lipofectamine 2000 (Life Technologies) according to the manufacturer’s manual. Expression was performed for 5 days. Alternatively, at DIV10, cells were transduced with 2.5 x 10^8^ viral genome copies (GC). The virus particles were directly added to each individual well. At DIV14 the cells were either fixed or subjected to treatment. Antimycin A treatments were performed over night at 37°C. Antimycin A (f.c. 1 µM) was directly added to each well, which contained 400 µL medium. D-Methionine (D-Met) treatment was performed 1 h at 37°C. The medium of neurons was removed and replaced by minimal NB medium, supplemented with 5 mM D-Met. Mock samples were treated with the same volume of solvent solution (PBS). After treatments neurons were fixed and used for ICC.

### ICC

Fixation was performed by replacing culture medium by 4% PFA in PBS. After incubation for 20 min at room temperature (rt), PFA was removed and cells washed twice with 50 mM NH_4_Cl in PBS, each time 10 min at rt. Subsequently, cells were blocked for 1 h at rt with blocking solution (10% goat serum, 1% BSA, 0.2% TritonX100, in PBS). After washing cells with PBS for 5 min at rt, antibody incubation was performed for 1 h at rt. Antibodies were diluted in PBS. Afterwards, three washing steps with PBS were performed, each 10 min at rt. Lastly, coverslips were mounted with Mowiol/Dabco and dried over night at rt. We used the following antibodies: anti-vesicular GABA transporter (vGAT) (1:1 000, #131003, *SYSY*); anti-GABA_A_Rγ2 (1:500, #224004, *SYSY*); anti-GephyrinE (3B11; 1:10, self-made); goat anti-mouse AlexaFluor 488 (1:500, #A-11029, *Invitrogen*), goat anti-rabbit AlexaFluor 647 (1:500, #A-21245, *Invitrogen*), goat anti-rabbit AlexaFluor 488 (1:500, #A-11075, *Invitrogen*).

### Confocal microscopy and image analysis

Pictures were tacking using the *Leica* TCS SP8 LIGHNING upright confocal microscope with an HC PL APO CS2 63x/1.30 glycerol objective. The microscope was equipped with hybrid *Leica* HyD detectors and diode lasers with 405, 488, 522 and 638 nm. The LIGHTNING adaptive deconvolution of the mounting medium “Mowiol” was used, which is capable of theoretical resolutions down to 120 nm and 200 nm lateral and axial, respectively. Images were acquired with stacks of 3 µm z-step size, 2048 x 2048 (144.77 x 144.77 µm) and segmented and analyzed in an automated fashion using ImageJ/FIJI 1.53t. Therefore, the previously described macros^57^ were used as a blueprint and adapted.

### Perfusion of mice and sample preparation

Mice were anesthetized with Ketamine/Xylazine through intra-peritoneal injection. After surgical tolerance was achieved, the abdomen was opened and liver and heart uncovered. A canula was pricked into the bottom of the left heart chamber to inject PBS containing 50 mM NEM into the cardiovascular system. A drain was given by a cut above the liver. PBS was injected until the liver lost the red color and turned pale. After decapitation, liver, brain and spinal cord were removed and added to RIPA buffer (50 mM Tris/HCl pH 8.0, 150 mM NaCl, 0.1% SDS, 1% IGEPAL CA-630, 0.5% Na deoxycholate, 1 mM EDTA, freshly added protease inhibitor (*Roche*)) containing 50 mM NEM. The tissue was dounce homogenized with 1 200 rpm 20 times, sonicated 10 sec at 4°C and 30% amplitude and subsequently centrifuged for 10 min at 4°C and 4 000 x g. The supernatant was transferred into a new tube and incubated for 1 h to let free thiols react with NEM. Afterwards, protein concentration was determined using BCA. Typically, 45 µg protein were used for SDS-PAGE for WB analysis.

### SDS-PAGE, Coomassie and Western Blot

Samples were supplemented with 1x sample buffer (5x: 250 mM Tris/HCl pH 6.8, 30% glycerol, 0.1% Bromophenol blue, 10% SDS) and incubated for 5 min at 95-98°C. We used either 5 mM β-ME for recombinant protein samples or 5 mM DTT for tissue samples or omitted it. Separation was performed with 5-12% SDS acrylamide gels using running buffer (3.029 g/L Tris, 14.4 g/L glycine, 0.1% SDS). Afterwards gels were either stained with coomassie (30% EtOH, 10% acetic acid, 0.25% Coomassie brilliant blue R250) or used for western blot. Therefore, the transfer was performed semi-dry using PVDF membranes and transfer buffer (3 g/L Tris, 24.15 g/L glycine pH 8.8, 10% MeOH). Blocking was performed for 1 h at rt under shaking in blocking solution (10% BSA in TBST). Primary and secondary antibodies (diluted in 1% BSA in TBST) were applied for 1 h at rt under shaking, with a wash step of 3 x 5 min with TBST (6.5 g/L Tris, 8.75 g/L NaCl pH 7.4, 0.05% Tween20) between primary and secondary antibody. After additional washing 3 x 5 min with TBST, detection was performed using ECL and the ChemiDocTM Imaging System (*BioRad*). The following antibodies were used: anti-GephyrinE (3B11; 1:10, self-made); anti-GAPDH (1:1 000, G9545, *Sigma*), anti-mouse HRP-coupled (1:10 000, AP181P, *Sigma*), anti-rabbit HRP-coupled (1:10 000, AP187P, *Sigma*)

### Statistics

The visualization and statistical analysis were performed with GraphPad Prism6 and R vers. 4.2.1. Standard deviation, individual data points and used tests are shown in the individual graph and in the captions respectively. Data points were identified as outliers when a two-fold standard deviation was exceeded. If data was normalized, *student’s* t-test was performed, and the significance level alpha corrected by the number of tests. Otherwise, we performed 1-way ANOVA and *Bonferroni* post-hoc tests. We analyzed: *N* = 6 individual mice for redox-dependent multimers; *N* = 3 technical replicates in the Moco activity assay, and *N* = 3 biological replicates; *N* = 3 biological replicates in SEC experiments; *N* = 3 biological replicates of gephyrin in ITC experiments; *N* = 3 biological replicates of gephyrin in co-sedimentation assays, *N* = 3 biological replicates of gephyrin in calpain cleavage assays; *N* = 10 cells per condition from 3 individual embryos of one neuron preparation in all experiments using hippocampal neurons.

## ACKNOWLEDGEMENTS

Technical help from Julia Reich is greatly appreciated. Furthermore, we thank Franziska Neuser and the Center for Mouse Genetics (CMG) of the University of Cologne for the organization of mice experiments. This work was financed by the german research foundation (DFG) (RTG2550 reloc to JR).

## AUTHOR CONTRIBUTIONS

MTG performed the experiments; MTG analyzed the data; MTG and GS wrote the manuscript; MTG, GS, JR designed the study; FL and LJ gave methodical input.

