## Supplementary Figures for "Redox-dependent synaptic clustering of gephyrin"

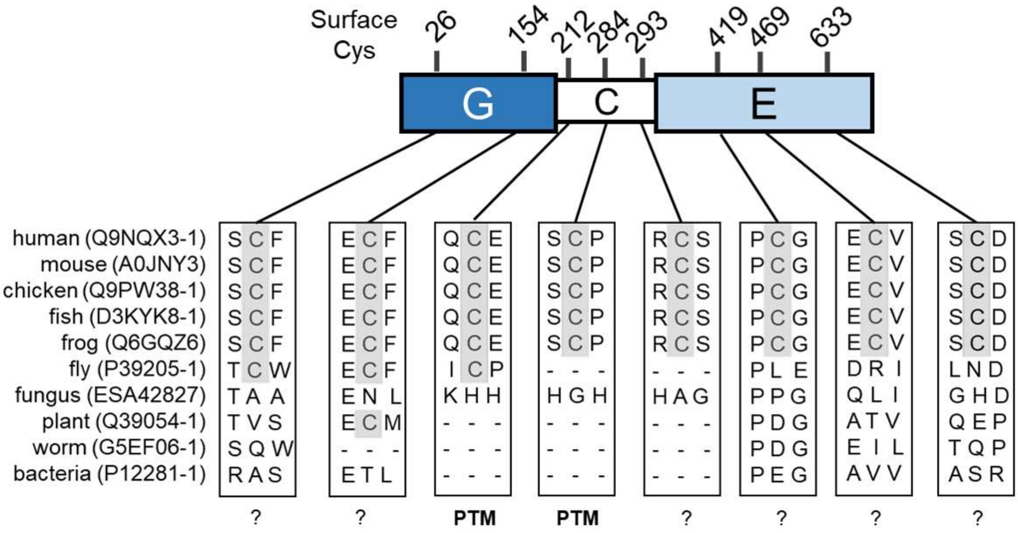


Figure S1: Alignment of Gephyrin and Homologues.

Alignment of protein sequence of different species. Areas around the chosen cysteines of gephyrin depicted with boxes. Cys 212 and 284 have assigned functions (targets of palmitoylation), while the function of the others is unknown (?).

---

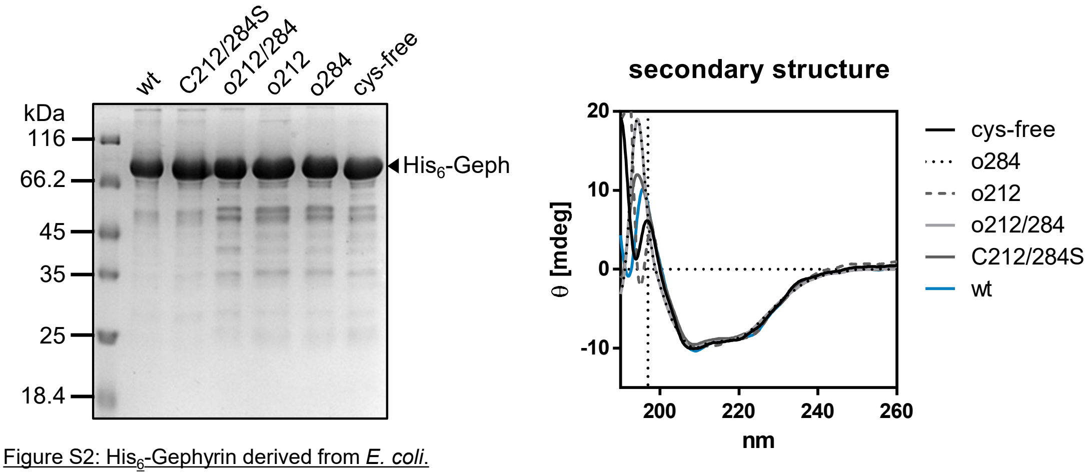


Figure S2: $His_6$-Gephyrin derived from *E. coli*.

$His_6$-Gephyrin variants recombinantly expressed and purified from *E. coli*. A) Coomassie gel of purified proteins. B) Analysis of secondary structure by CD Spec. Verical dotted line indicates data points with HT values above 800 mV (machine threshold).

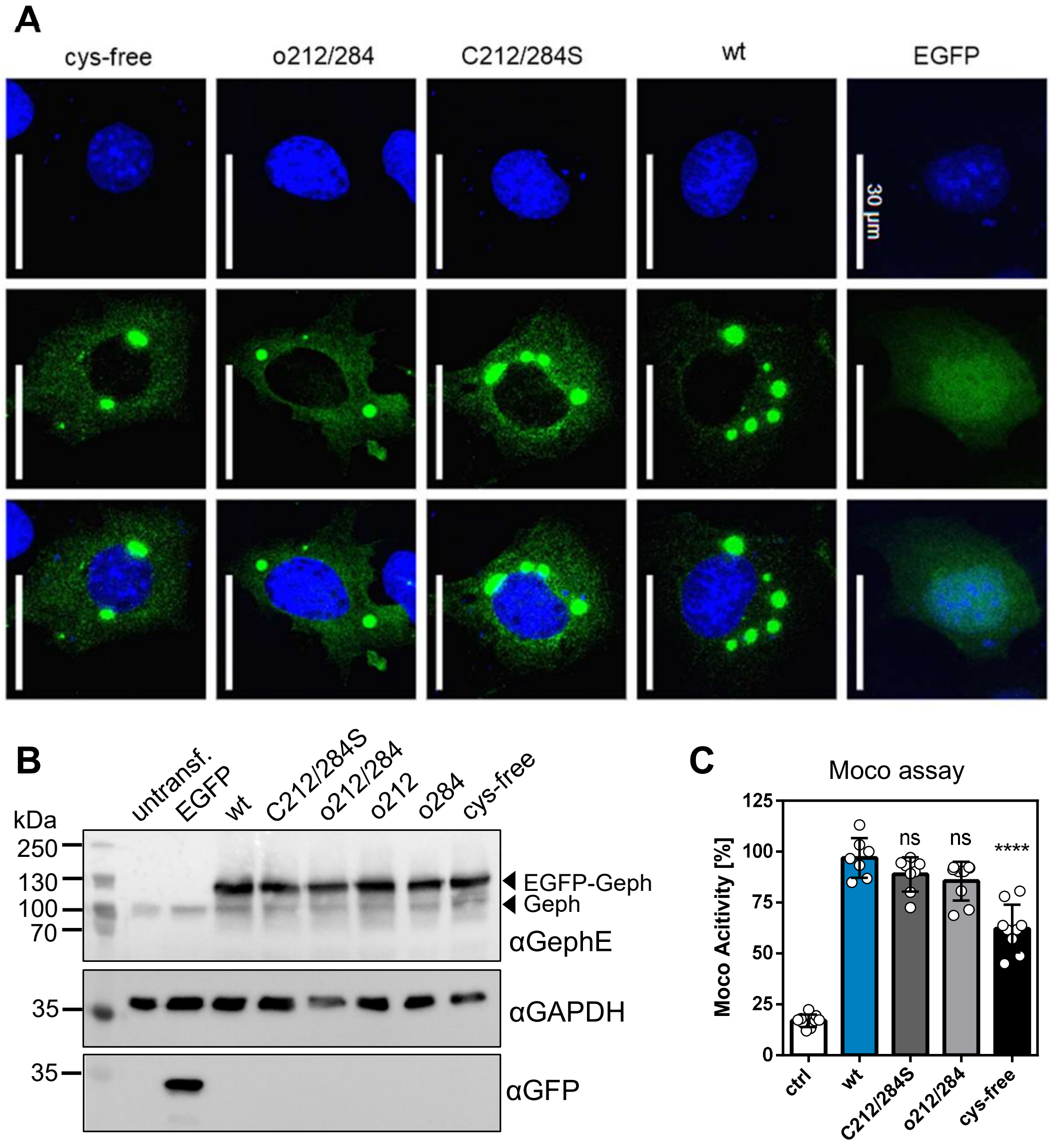


Figure S3: Expression of EGFP-Geph in COS7 and HEK293 Cells.

EGFP-Gephyrin variants were recombinantly expressed in COS7 and HEK293 cells for 48 h. A) Images of COS7 cells. Blue: DAPI, green: EGFP-Geph, scale bar of 30 µm. B) Expression in HEK293 cells. Protein amounts analyzed by Western blot. EGFP-Geph and endogenous gephyrin marked with arrows. C) Moco assay with cysteine variants. Moco activity normalized to wt. Significance towards wt tested by student's t-test, alpha level corrected by three. C212/284S: $p$=0.3288 (ns); o212/284: $p$=0.1026 (ns); cys-free: $p$<0.0001 (****).

---

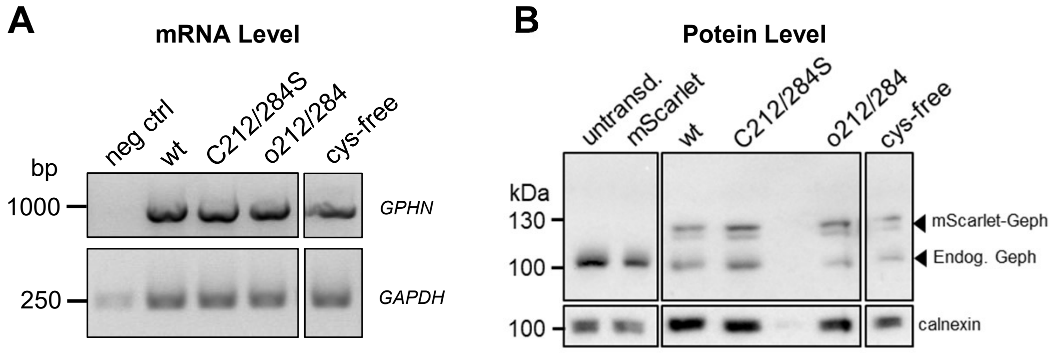


Figure S4: Expression of recombinant Gephyrin in Hippocampal Neurons.

Expression of mScarlet-gephyrin via transduction in primary hippocampal neurons. A) mRNA level determination. Isolated mRNA of mSclt-Geph was used for PCR. Primers were designed to exclusively bind to recombinant gephyrin. GAPDH used as a ctrl. B) Western Blot of neuronal lysates. Endogenous and recombinant gephyrin species indicated by arrows.

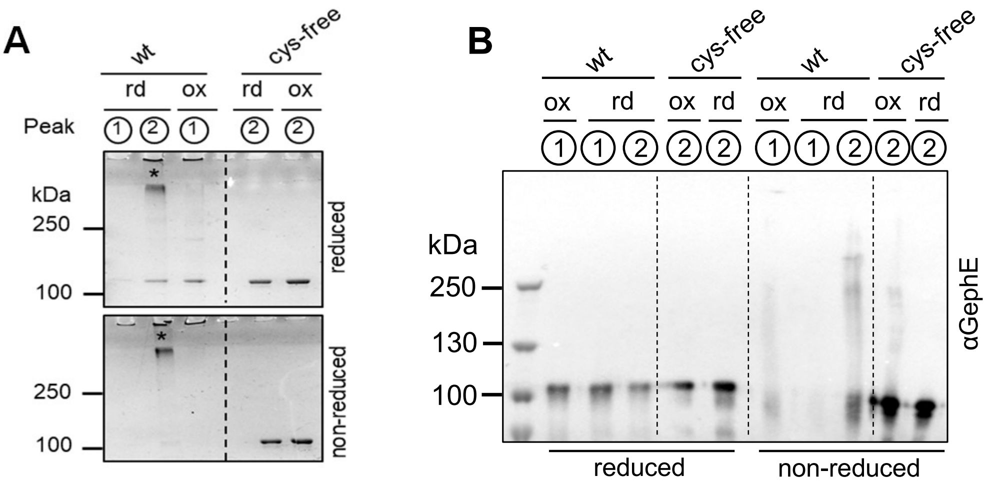


**C**

| Variant | $N^{high}$ | ± | $K_D^{high}$ [µM] | ± | $H^{high}$ [kcal/mol] | ± | $S^{high}$ [cal/mol/deg] |
|---|---|---|---|---|---|---|---|
| wt_1 | 0.302 | 0.006 | 0.0123 | 0.0029 | -28.47 | 0.00 | -55.60 |
| cysfree_1 | 0.438 | 0.016 | 0.0763 | 0.0238 | -31.53 | 0.72 | -69.10 |
| wt_2 | 0.360 | 0.005 | 0.0249 | 0.0074 | -31.01 | 0.54 | -65.20 |
| cysfree_2 | 0.443 | 0.011 | 0.0391 | 0.0203 | -29.34 | 0.74 | -60.70 |
| wt_3 | 0.349 | 0.006 | 0.0870 | 0.0191 | -63.31 | 1.58 | -172.00 |
| cysfree_3 | 0.342 | 0.006 | 0.0735 | 0.0017 | -65.20 | 1.64 | -178.00 |

| Variant | $N^{low}$ | ± | $K_D^{low}$[µM] | ± | $H^{low}$ [cal/mol] | ± | $S^{low}$ [cal/mol/deg] |
|---|---|---|---|---|---|---|---|
| wt_1 | 0.492 | 0.012 | 0.877 | 0.1808 | -11.82 | 0.65 | -10.40 |
| cysfree_1 | 0.516 | 0.036 | 1.366 | 0.3919 | -8.10 | 2.01 | 0.71 |
| wt_2 | 0.920 | 0.027 | 0.787 | 0.2530 | -5.40 | 0.43 | 10.50 |
| cysfree_2 | 0.880 | 0.049 | 0.971 | 0.5316 | -4.76 | 0.83 | 12.20 |
| wt_3 | 0.952 | 0.031 | 1.412 | 0.3471 | -10.93 | 1.13 | -8.48 |
| cysfree_3 | 0.964 | 0.030 | 1.259 | 0.3283 | -10.69 | -0.01 | -7.49 |

**D**

| Variant | $N^{high}$ | ± | $K_A^{high}$ [$M^{-1}$] | ± | $H^{high}$ [kcal/mol] | ± | $S^{high}$ [cal/mol/deg] |
|---|---|---|---|---|---|---|---|
| wt_rd_1 | 0.716 | 0.007 | 0.226 | 0.030 | -12.01 | 0.19 | -11.20 |
| wt_ox_1 | 0.728 | 0.006 | 0.281 | 0.053 | -11.05 | 0.16 | -7.56 |
| cys-free_rd_1 | 0.739 | 0.001 | 0.203 | 0.035 | -10.94 | 0.23 | -7.84 |
| cys-free_ox_1 | 0.750 | 0.005 | 0.429 | 0.056 | -10.06 | 0.13 | -3.40 |
| wt_rd_2 | 0.687 | 0.007 | 0.467 | 0.059 | -12.44 | 0.19 | -12.80 |
| wt_ox_2 | 0.729 | 0.005 | 0.408 | 0.043 | -11.02 | 0.13 | -7.72 |
| cys-free_rd_2 | 0.778 | 0.005 | 0.714 | 0.059 | -12.74 | 0.13 | -14.60 |
| cys-free_ox_2 | 0.750 | 0.004 | 0.261 | 0.022 | -9.88 | 0.08 | -3.02 |
| wt_rd_3 | 0.712 | 0.006 | 0.267 | 0.038 | -8.41 | 0.12 | 1.86 |
| wt_ox_3 | 0.798 | 0.006 | 0.195 | 0.028 | -8.06 | 0.11 | 3.68 |
| cys-free_rd_3 | 0.809 | 0.006 | 0.201 | 0.029 | -7.61 | 0.09 | 5.13 |
| cys-free_ox_3 | 0.769 | 0.012 | 0.183 | 0.019 | -8.39 | 0.24 | 1.10 |

Figure S5: Analysis of SEC fractions and ITC Data

A and B: Fractions taken of Peak 1 (high-order multimers) and Peak 2 (hexamers) of the respective runs using reduced (rd) or oxidized gephyrin wt or cys-free variant. The samples have been prepared for SDS-PAGE with a reducing or a non-reducing buffer. A) Coomassie Gel. Unspecific band marked with a star (*). B) Western blot using antibody against Gephyrin E-domain. C and D: Interaction parameters of gephyrin wt or cys-free variant towards different ligands determined by mathematical fitting of ITC runs. A) ITC experiment titrating GlyRβICD peptide into gephyrin under reductive conditions. Data of two-site fit. B) ITC experiment titrating LSβICD into pre-reduced or -oxidzed gephyrin. Data of one-site fit.

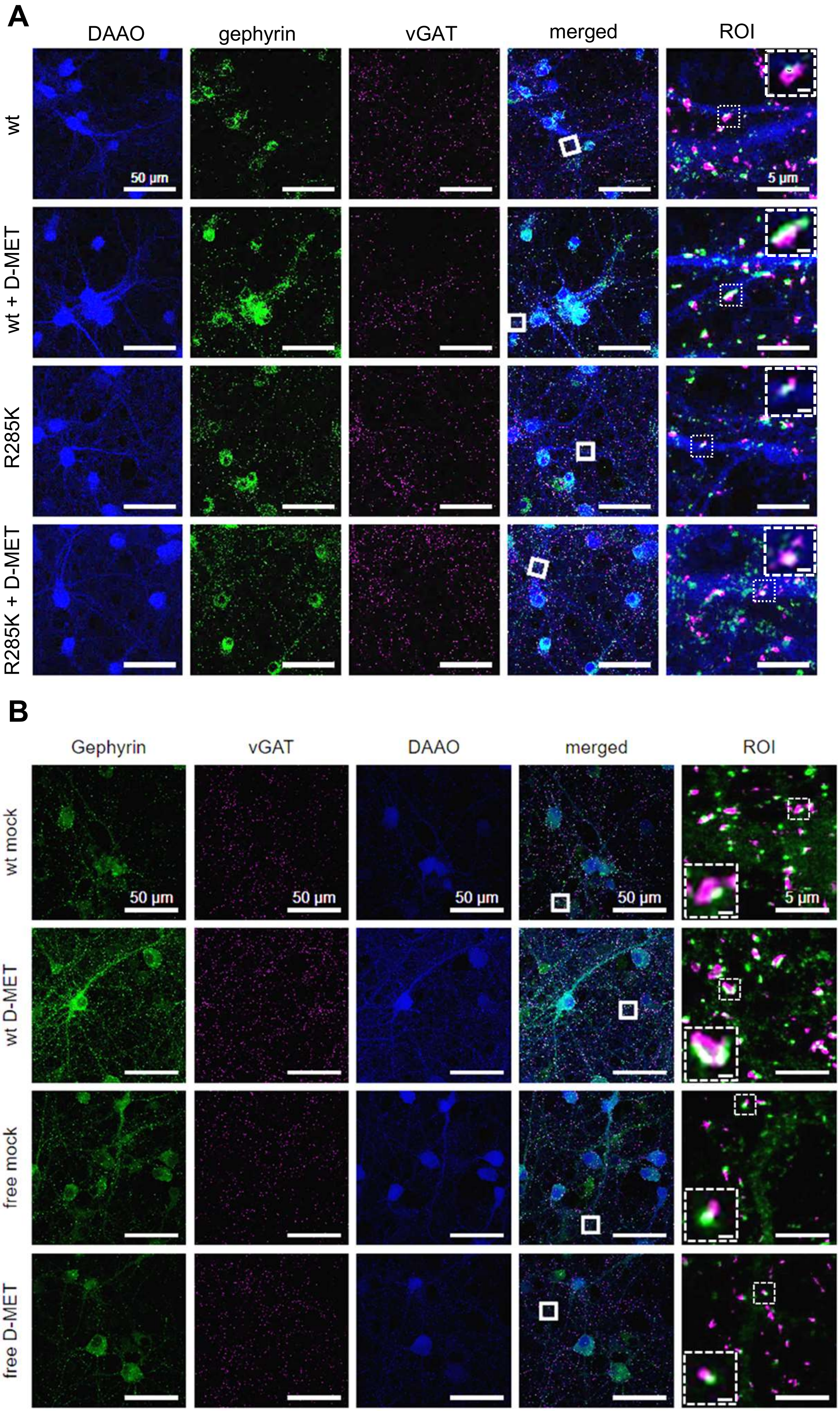


Figure S6:$H_2O_2$ drives Gephyrin Oligomerization at Synapses.

Expression of cytosolic mSclt-DAAO and mEGFP-Geph in primary, hippocampal neurons. Treatment at DIV14 with 5 mM D-MET. Analysis of cluster sizes [$\mu m^2$], intensity [au] and co-localization with vGAT (=synaptic clusters) of gephyrin. A) mSclt-DAAO wt and R285K. Analysis of endogenous gephyrin clusters. Gephyrin: green, vGAT: magenta, DAAO: blue; scale bar 30 and 0.5 µm. B) mSclt-DAAO wt and mEGFP-Geph wt and cys-free variants. Analysis of recombinant mEGFP-gephyrin clusters. Gephyrin: green, vGAT: magenta; scale bar 30 and 0.5 µm.

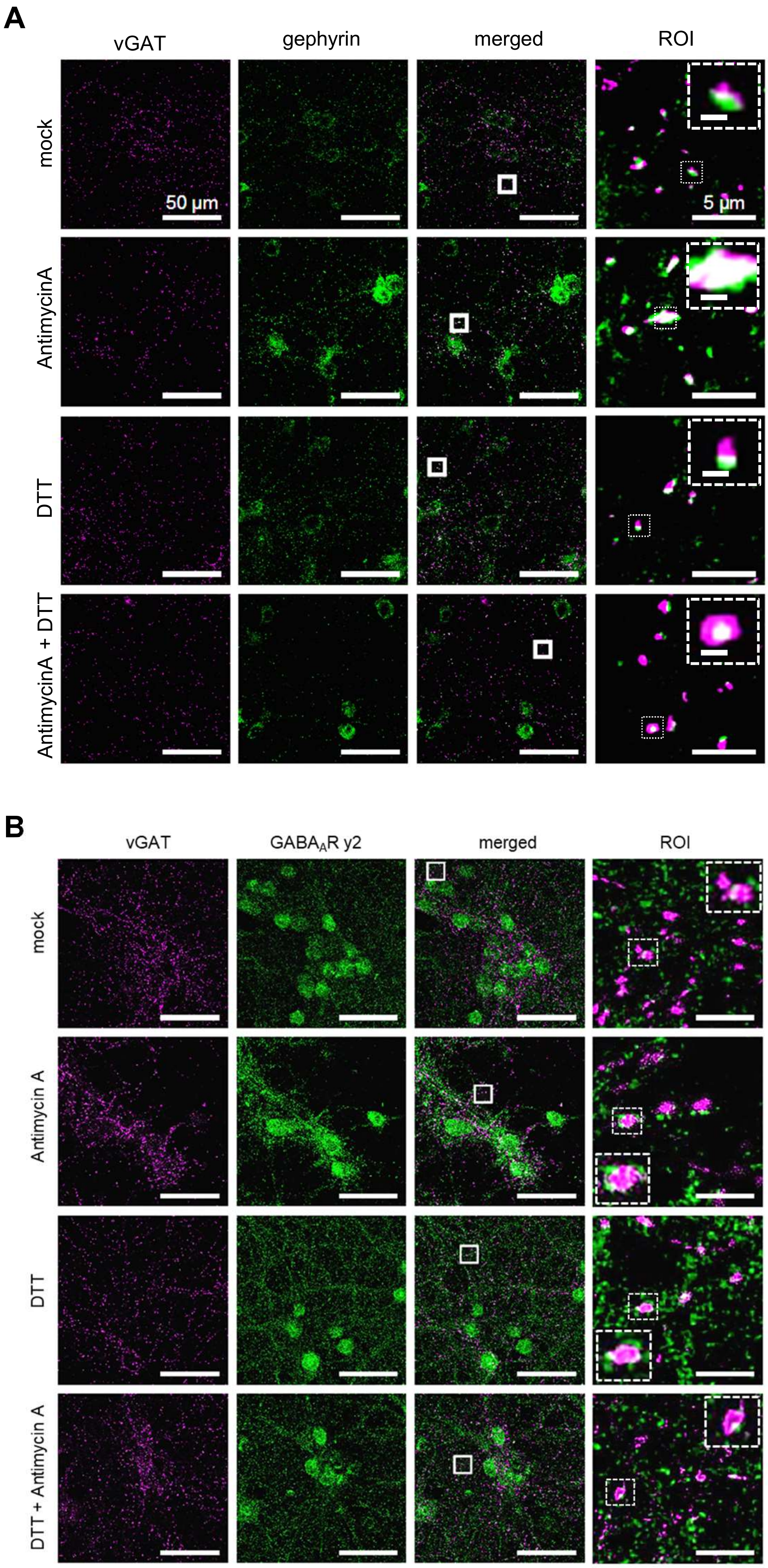


Figure S7: mROS reversibly drive Gephyrin Oligomerization and the Recruitment of $GABA_AR$ γ2 Subunits

Treatment of primary hippocampal neurons at DIV14 with 1 µM Antimycin A (AntiA) and/or 5 µM DTT. Analysis of gephyrin or $GABA_AR$ γ2 subunit cluster sizes [$\mu m^2$], intensity [au] and co-localization with vGAT (=synaptic clusters). A) Analysis of endogenous gephyrin. Gephyrin: green, vGAT: magenta, DAAO: blue; scale bar 50 and 0.5 µm. B) Analysis of endogenous $GABA_AR$ γ2 subunit clusters. γ2: green, vGAT: magenta; scale bar 50 and 0.5 µm.
